# Otx2 and Oc1 directly regulate the transcriptional program of cone photoreceptor development

**DOI:** 10.1101/2020.06.29.177428

**Authors:** Nicolas Lonfat, Su Wang, ChangHee Lee, Jiho Choi, Peter J. Park, Connie Cepko

## Abstract

The vertebrate retina is a highly organized structure of approximately 110 cell types. Retinal progenitor cells (RPCs) produce these cell types in a temporal order that is highly conserved. While some RPCs produce many cell types, some terminally dividing RPCs produce restricted types of daughter cells, such as a cone photoreceptor and a horizontal cell (HC). Here, we compared the transcriptomes and chromatin profiles of such a restricted cone/HC RPC with those of other RPCs. We identified many cis-regulatory modules (CRMs) active in cone/HC RPCs and developing cones. We then showed that Otx2 and Oc1 directly regulate the activity of multiple CRMs genome-wide, including near genes important for cone development, such as Rxrg and Neurod1. In addition, we found that Otx2 regulates itself. These results suggest that Otx2 and Oc1 have a broader role than previously appreciated, and deepen our understanding of retinal development, which may benefit therapies for retinal diseases.

## Introduction

During development, retinal progenitor cells (RPCs) divide to give rise to an extremely complex and highly organized tissue. The retina is composed of 6 major types of neurons that are born in a stereotypical and overlapping order, conserved across vertebrates ^1–5^. The retinal ganglion cells (RGCs) are born first, followed by the cone photoreceptors concomitant with the horizontal cells (HCs), followed by the majority of the amacrine cells (ACs). Rod photoreceptors and bipolar cells (BPs) are born late, along with a glial cell type, the Mueller Glial (MG) cell. How RPCs produce such a diversity is a question of interest, both for our understanding of the development of a complex tissue, and for therapeutic applications. Retinal diseases involving cones are common, with many genetic lesions leading to the dysfunction, and sometimes the death, of these photoreceptors ^6,7^. As cones are the type of photoreceptor that we use to initiate color and daylight vision, their loss can be devastating. Cone degeneration also leads to loss of high acuity vision, particularly in a disease that is growing in frequency, Age-Related Macular Degeneration (AMD) ^8^. One therapeutic approach is to culture stem cells or retinal organoids and induce them to produce cones, either for engraftment, or to serve as in vitro disease models ^9^. In addition, gene therapy vectors that express therapeutic genes only in cones are under development ^10^. A better understanding of the gene regulatory networks (GRNs) that lead to cone genesis and cone gene regulation may benefit such applications.

Many RPCs are multipotent, capable of giving rise to combinations of many types of retinal cells, as shown by lineage studies in several species ^11^. Even when clonal marking was initiated at a single time, clones had a highly variable composition and size ^12–14^. Moreover, although birthdating data have shown a conserved order in genesis of different cell types, there is overlap, in that RPCs can make different cell types at one time ^1–5^. These observations have raised the question of whether all RPCs are equivalent ^15^. In part to address this question, a comprehensive cataloguing of molecular differences among RPCs was initiated by RNA profiling of single RPCs using microarrays ^16^, and more recently, by RNA-seq ^17–19^. These data revealed many different combinations of genes expressed in RPCs, and inspired studies to ask whether differences in RPC gene expression correlated with production of different types of progeny. In mice, RPCs that express the bHLH transcription factor (TF), Olig2, were shown to be terminally dividing, producing only cones and HCs embryonically, and only rods and amacrine cells postnatally ^20^. Zebrafish cones and HCs also were shown to share a common lineage, through live imaging of RPCs expressing a reporter for the cone gene, Thyroid hormone receptor beta (Thrb) ^21^. Similarly, chick RPCs expressing a reporter driven by an enhancer of Thrb were biased to the production of HCs and photoreceptors ^22^.

We previously characterized a CRM for Thrb, designated CRM1, and discovered that it is regulated by the TFs Otx2 and Oc1 ^22^. We showed that Oc1 was not only a regulator of Thrb, but it contributed to the choice of cone vs rod fate. Genetic studies in mice showed that Onecut genes are required beyond photoreceptors, as Oc1 and Oc2 are involved in RGC and HC development ^23^. Null mutations of Otx2 showed that it is required for both rod and cone development in mice ^24^. Interestingly, Otx2 is also required for BP and HC fates ^24–26^, even though it is not expressed in HCs, and HCs do not share specific features with photoreceptors.

To further investigate the molecular mechanisms that restrict an RPC to the production of cones and HC, we used ThrbCRM1 as an entry point to this question. Genome-wide methods were used to examine the chromatin status and transcriptomes of ThrbCRM1+ RPCs vs those of ThrbCRM1-RPCs. Thousands of DNA sequences were differentially open in ThrbCRM1+ RPCs, with motifs for Otx2 and Oc1 showing the most enrichment within these regions. The activity of many of the predicted CRMs with these TF binding sites (TFBSs) was tested in retinal tissue, which showed them to be active in early cones and/or the cone/HC RPC. These CRMs included one near Neurod1, a gene previously shown as important in cone development, as a regulator of Thrb ^27^, and one near Rxrg, a known partner of Thrb, also involved in cone patterning ^28,29^. The CRM activity of most of these sequences was found to be dependent on the binding of both Otx2 and Oc1, and thousands of other predicted binding sites for Otx2 and Oc1 were validated via chromatin binding assays using antibodies to Otx2 and Oc1. In addition, we directly searched for the CRMs that regulate Otx2 and Oc1 and found that Otx2 regulates itself. These findings show that Otx2 and Oc1 coordinate the development of cones, via the direct regulation of multiple genes important for the development and function of this cell type.

## Results

### Integrating RNA expression and open chromatin to discover CRMs for cone development

In order to perform a genome-wide search for potential CRMs for developing cones, we used ATAC-seq to profile the open chromatin regions in early embryonic chick retinal tissue, from the peak period of cone genesis ^1,3^. To enrich for RPCs that were producing cones, and for newly postmitotic cells fated to be cones, we used GFP expression driven by ThrbCRM1 ^22^. This strategy was taken as electroporation tends to target mitotic cells and the plasmids are inherited and expressed by their newly post-mitotic progeny ^30^. Freshly explanted embryonic day 5 (E5) retinas were co-electroporated with the ThrbCRM1-GFP plasmid and the broadly expressed plasmid, CAG-mCherry. After 2 days in culture, the electroporated cells were FACS-sorted into ThrbCRM1+/CAG-mCherry+ (referred to hereafter as ThrbCRM1+) and ThrbCRM1-/CAG-mCherry+ (referred to hereafter as ThrbCRM1-) cell populations. We processed them for ATAC-seq to identify regions of chromatin that were differentially open in ThrbCRM1+ cells relative to those of ThrbCRM1− cells (Figure 1A). Profiles were highly reproducible among the three and two replicates for the ThrbCRM1+ and ThrbCRM1− cells, respectively, and we combined the replicates for the comparison of their chromatin accessibility.

**Figure 1:**
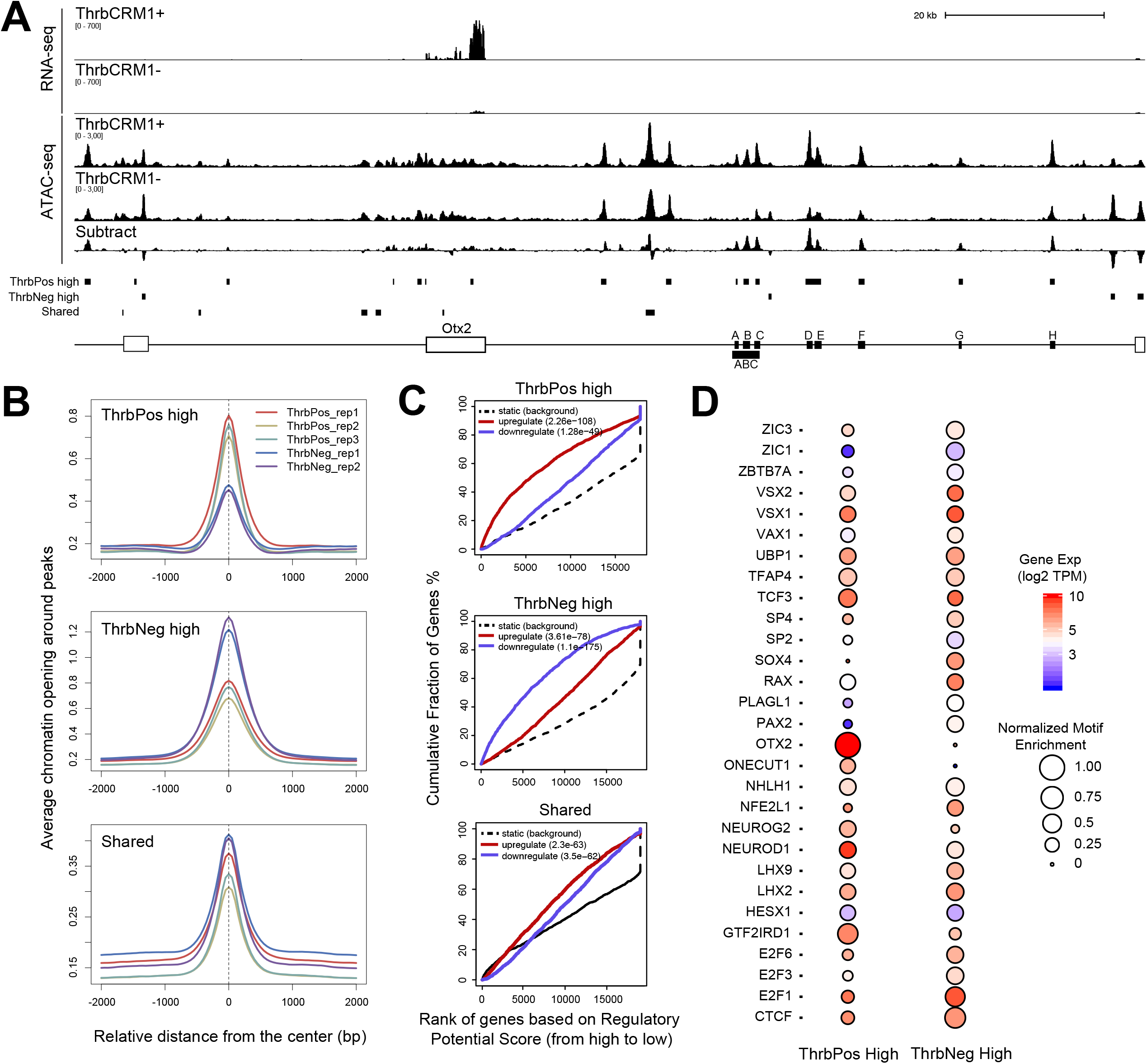
Integrating RNA expression and open chromatin for discovery of CRMs. ATAC-seq data from ThrbCRM1+ and ThrbCRM1− cells from the chick retina were integrated with the corresponding RNA-seq. (A) The Otx2 locus is shown, with the RNA-seq data on top, followed by the ATAC-seq and the corresponding differential track created by subtraction of the ThrbCRM1− cell peaks from the ThrbCRM1+ cell peaks. ATAC-seq peaks called for ThrbCRM1+ cells (ThrbPos high), ThrbCRM1− cells (ThrbNeg high) or that were shared between conditions are shown as small black rectangles. The regions A-H (black rectangles) which were tested for CRM activity (Figure 2) are shown near the Otx2 gene (white boxes). (B) The average chromatin opening across ThrbPos high peaks (top), ThrbNeg high peaks (middle) and peaks shared between conditions (bottom) in different replicates. (C) BETA activating/repressive function prediction of the ThrbPos high, ThrbNeg high and shared ATAC-seq peaks in the two ThrbCRM cell populations. The colored lines represent upregulated and downregulated genes in the indicated population of cells. The dashed line indicates the non-differentially expressed genes as background. Genes are accumulated by their rank on the basis of the regulatory potential score from high to low. The regulatory potential shown on the x-axis represents the likelihood of a gene being more highly expressed in the indicated population of cells, calculated by considering both the accessibility and the distance between the gene TSS and the ATAC-seq peak (see details in Methods). P-values that represent the significance of the upregulated or downregulated gene group distributions are compared with the static gene group by the Kolmogorov–Smirnov test. (D) Comparison of motif enrichment in regions open in ThrbCRM1+ and ThrbCRM1− cells. Size of the circle represents the significance of the motif. Motifs that were not expressed at least in one population were excluded.

We found ~96,500 and ~103,000 peaks of open chromatin for ThrbCRM1+ and ThrbCRM1− cells, respectively (Sup Tables 1). Overall, open chromatin profiles were highly concordant between ThrbCRM1+ and ThrbCRM1− cells (example for Otx2 shown in Figure 1A). To identify the regions with differential chromatin accessibility throughout the genome, we subtracted the aligned ATAC-seq peak reads of ThrbCRM1− from those of ThrbCRM1+ to generate a difference profile (Figure 1A). Peaks were then called using this profile to nominate regions either enriched in ThrbCRM1+ or ThrbCRM1− cells (Sup Tables 1). We intersected these peaks with the original sets of peaks for each condition, to identify them as ThrbPos high, ThrbNeg high or shared peaks (Figure 1A,B; Sup Tables 1). Although ThrbPos high peaks and ThrbNeg high peaks were more open in ThrbCRM1+ and ThrbCRM1− cells, respectively, these regions tended to be accessible in both populations (Figure 1B).

To further characterize the DNA sequences potentially relevant for cone development, we compared these ATAC-seq profiles with the RNA-seq data available from the same cell populations ^31^ (Figure 1A, Sup Tables 2). The Binding and Expression Target Analysis (BETA) ^32^ was used to examine the expression of genes within a window of 100 Kb of an ATAC-seq peak (Figure 1C, Sup Tables 2). Overall, ThrbPos high peaks were more likely to be open around genes that were upregulated in ThrbCRM1+ cells relative to ThrbCRM1− cells. Conversely, ThrbNeg high peaks were associated with genes that were upregulated in ThrbCRM1− cells. The shared peaks were open around genes that were expressed in both populations at similar levels (Figure 1C, Sup Tables 2). This analysis resulted in a similar trend using a larger window of 500kb and 1Mb (not shown).

We then searched for enriched TFBS motifs in differential peaks that might contribute to the regulation of the differentially expressed genes. We identified motifs predicted in both ThrbPos high and ThrbNeg high enriched peaks (Sup Tables 3), and filtered out the cognate TFs that were not expressed in these cells (Figure 1D). Only a few predicted motifs were differentially enriched in each population. Interestingly, motifs for Otx2 and Oc1, the two TFs regulating Thrb via the ThrbCRM1 enhancer, were among the most differential predicted TFBSs, enriched strongly within ThrbPos high peaks (Figure 1D). Although the proneural TF, Neurog2, was expressed at similar levels in both ThrbCRM1+ and ThrbCRM1− cells, the Neurog2 binding site was more enriched in ThrbPos high peaks. A predicted motif for the TF Gtf2ird1, more highly expressed in ThrbCRM1+ cells, also was found more often in ThrbCRM1+ enriched peaks compared to those from ThrbCRM1− cells. Gtf2ird1 has been shown to modulate photoreceptor gene expression, to pattern cone opsin expression in cooperation with Thrb, and to be critical for photoreceptor function ^33^. A motif for the TF Neurod1, another important gene in cone development ^27^ and highly up-regulated in ThrbCRM1+ cells, was found in both ThrbPos and ThrbNeg high peaks (Figure 1D). TFBSs that were enriched in ThrbCRM1− cells, included motifs for Zic1, Zic3, Sp2, Sp4, Pax2, Plagl1 or E2f cell cycle genes. Surprisingly, the Sox4 motif was almost absent in ThrbCRM1+ cells, although the gene showed a higher expression level in those cells. Motifs for the progenitor genes, Vsx2 and Rax, which are expressed at a higher level in ThrbCRM1− cells, were predicted in both cell populations (Figure 1D).

### Identification of Otx2 and Oc1 regulatory elements using ATAC-seq

Multiple differential ATAC-seq peaks were found around genes more highly expressed in the ThrbCRM1+ cells (Figure 1C). As we are interested in the networks that might regulate cone development, and Otx2 and Oc1 are two genes important in the early stages of cone development, we first compared the open chromatin profiles of the ThrbCRM1+ and ThrbCRM1− cells at the Otx2 and Oc1 loci. We selected peaks of open chromatin near Otx2 that were specifically enriched in ThrbCRM1+ cells (Figure 1A). To test if these regions indeed have regulatory activities, we cloned the corresponding regions into the Stagia3 reporter plasmid driving GFP and placental alkaline phosphatase (PLAP) ^34^. The plasmids were electroporated along with a co-electroporation control (CAG-mCherry) into E5 chick retinas, which were then cultured for 2 days. Alkaline phosphatase (AP) staining was performed to assess the activity of the potential CRMs. Two CRMs near the Otx2 gene induced strong AP signal (Figure 2A) and were designated Otx2 CRM E and F (Figure 1A, 2A).

**Figure 2:**
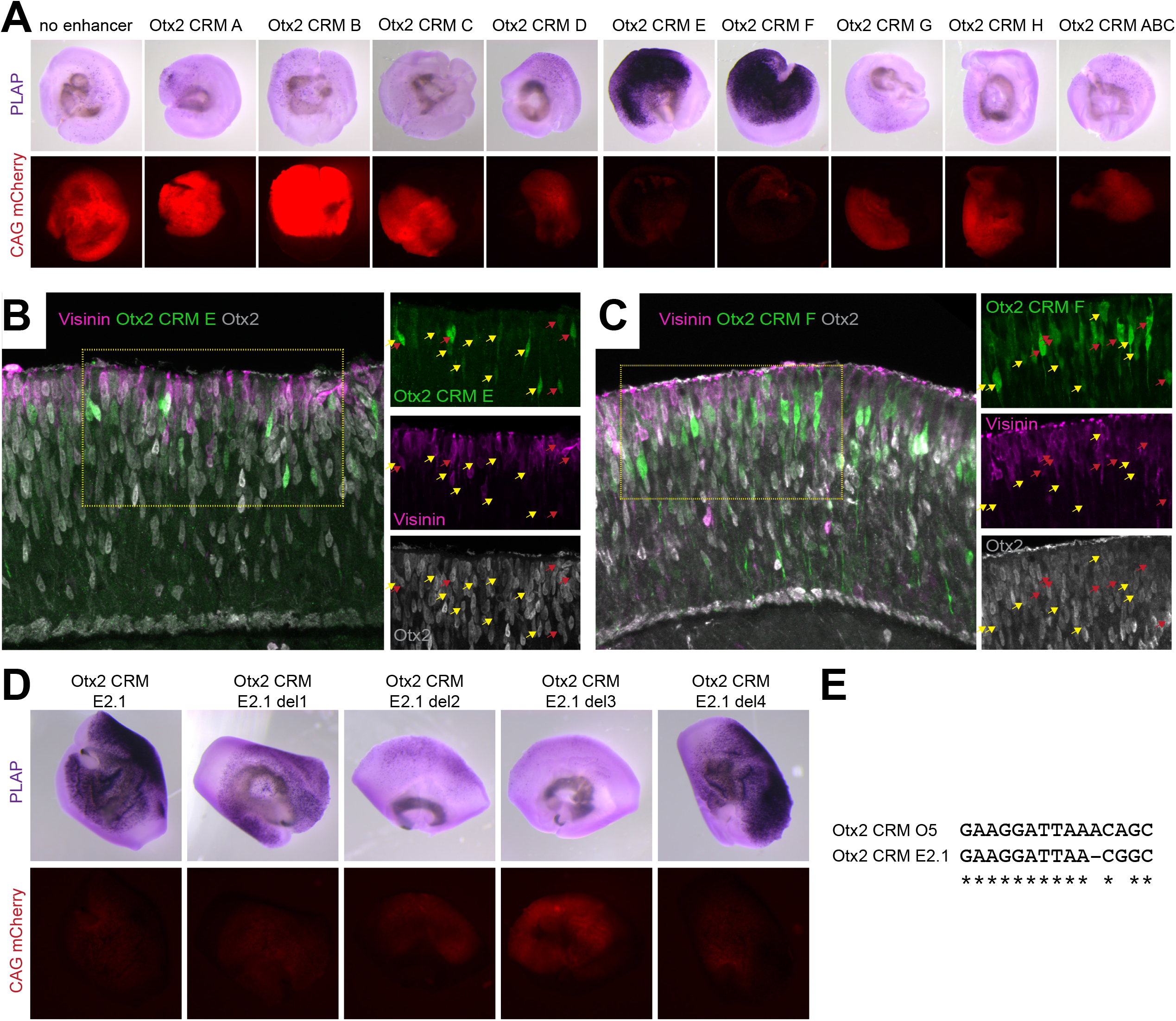
Assay of Otx2 regulatory elements predicted by ATAC-seq. (A) Potential Otx2 CRM A-H (Figure 1A) were assayed for activity by plasmid electroporation in chick retinal explants from E5-E7. Stagia3 plasmid encoding AP and GFP was used to assay for CRM activity (purple stain), and was co-electroporated with a control plasmid (CAG-mCherry, red). Corresponding images of each electroporated explant are shown (note that the AP reaction product absorbs the red fluorescent signal). (B-C) Sections of chick retinal explants (E5-E7) electroporated with Otx2 CRM E and CRM F are shown using the Stagia3 GFP readout. IHC for Otx2 protein (gray) or visinin (purple), a photoreceptor marker, are shown. Examples of cells with GFP expression that also expressed Otx2 protein are shown with yellow and red arrows, with those that also showed visinin indicated with red arrows. (D) Deletion analysis of Otx2 CRM E was used to identify the critical nucleotides for activity. The fragment Otx2 CRM E2.1 was found to be necessary and was searched for potential TFBSs using TRANSFAC. Deletions of 4 clusters of TFBSs (del1-4) within Otx2 CRM E2.1 showed that 3 regions were necessary. (E) The sequence deleted in del3 construct had a predicted TFBS for Otx2. The del3 sequence is conserved with the mouse Otx2 O5 CRM that requires Otx2 TFBS for activity ^40^.

To assess CRM activity at cellular resolution, GFP expression was examined in tissue sections of electroporated retinas. Otx2 CRM E and F were active in cells on the apical side of the retina, with the morphology and location of developing photoreceptor cells. A fraction of the cells also were positive for visinin (Figure 2B,C), a marker of photoreceptors ^35,36^ with a slightly later onset of expression than Thrb ^37^. GFP expression correlated with the presence of the Otx2 protein, detected using immunohistochemistry (IHC) (Figure 2B,C). These observations are consistent with these CRMs acting as enhancers for the Otx2 gene within RPCs and developing photoreceptor cells. In addition, Otx2 CRM F showed expression in more basal cells, which are potentially early HCs (Figure 2C), localized basally at this time during development ^38^.

Using deletion analysis, we refined the minimal DNA sequence necessary for Otx2 CRM E and F activity (Figure S1A). A 161bp sequence for region E (named Otx2 CRM E2.1), and a sequence of 410 bp for region F (Otx2 CRM F2), were able to drive AP activity (Figure 2D, S1A,B). The TRANSFAC prediction algorithm was used to search for potential TFBSs within the Otx2 CRM E2.1 sequence. Four clusters of TFBSs were identified, which were deleted and tested in the reporter assay (Figure S1C). The deletion of 3 out of the 4 regions resulted in a significant decrease in expression compared to the wild-type (WT) sequence (Figure 2D). One of the regions had a predicted TFBS for Otx2, which lost almost all expression upon deletion (del3) (Figure 2D). Interestingly, we recently found an enhancer for the murine Otx2 gene, named Otx2 O5, which is active in developing cones ^39^, as well as in mature BPs ^40^. In BPs, it showed a similar auto-regulation by Otx2 ^40^. We aligned the chick Otx2 CRM E2.1 sequence to the mouse Otx2 O5 sequence and found that the critical TFBS deleted in Otx2 CRM E2.1 (del3) was conserved with the TFBS critical for the mouse Otx2 O5 activity (Figure 2E).

Using predictions from the ATAC-seq peaks, we also identified regions with CRM activity at the Oc1 locus (Figure 3A). Multiple peaks were tested (n=13), with 9 of them showing AP activity (Figure 3B, S2A). While Oc1 CRM B, J, K and L have not been previously described as potential CRMs, the other CRMs were recently reported to have activity in the chick retina ^41^. The activity of the regions more open in ThrbCRM1+ cells that showed the strongest AP staining was then examined using GFP expression within tissue sections (Figure 3C). Oc1 CRM A and L positive cells were mostly found within the basal region of the retina, where HCs are found at this stage ^38^. The activity of the Oc1 CRM A was usually seen in cells expressing the Oc1 protein, and the Lhx1 protein, a marker of HCs ^42^, while expression from the Oc1 CRM L colocalized mostly with Oc1 only. A minority of cells positive for these two Oc1 CRMs had the morphology of early photoreceptors (Figure 3C). Oc1 CRM B, J and K were active in cells found in the apical region, also resembling photoreceptors. In these cells, we could not detect Oc1 protein (Figure 3C), consistent with a previous report that the gene is turned off as cells leave the RPC state and become photoreceptors ^43^. We also looked at the cellular activity of the Oc1 CRM G, which showed a similar chromatin accessibility between ThrbCRM1+ and ThrbCRM1− cells. As described recently for an overlapping CRM, Oc1 CRM ACR8 ^41^, the activity of Oc1 CRM G activity was found excluded from ThrbCRM1+ cells in what could be other types of RPCs (Figure S2B). Additionally, we found higher expression driven by Oc1 CRM G in cells morphologically resembling RGCs (Figure S2B). Taken together, this set of CRMs, nominated by ATAC-seq peaks, revealed potential regulatory elements for Otx2 and Oc1, operating at overlapping, early stages during the development of cones and HCs.

**Figure 3:**
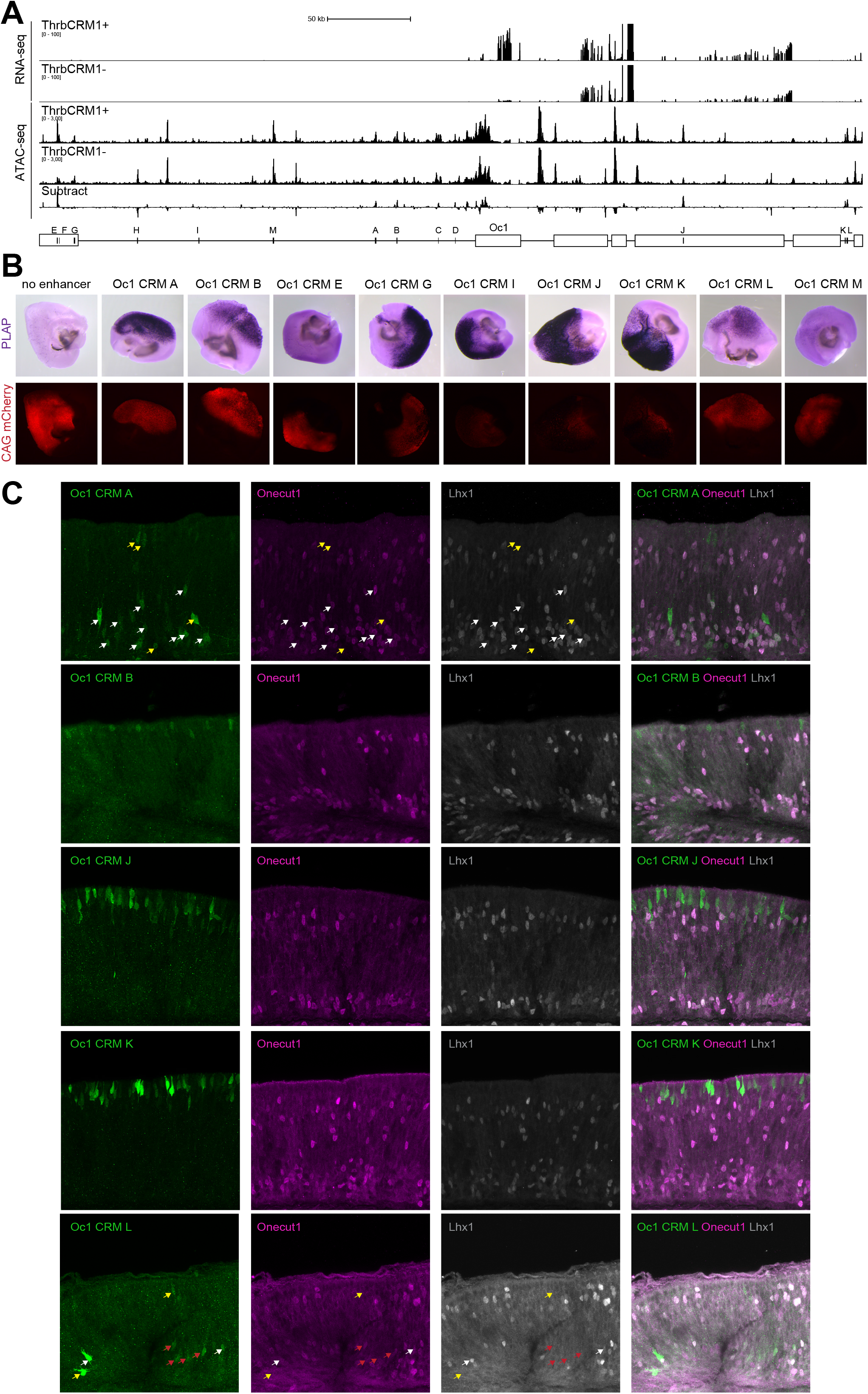
Assay of Oc1 regulatory elements predicted by ATAC-seq. (A) ATAC-seq data from ThrbCRM1+ and ThrbCRM1− cells from the chick retina were integrated with the corresponding RNA-seq. The Oc1 locus is shown, with the RNA-seq data on top, followed by the ATAC-seq data and the corresponding differential track created by subtraction of the ThrbCRM1− cell peaks from the ThrbCRM1+ cell peaks. The regions A-M (black rectangles) which were tested for CRM activity are shown near the Oc1 gene (white boxes). (B) Potential Oc1 CRM A-M were assayed for activity by plasmid electroporation in chick retinal explants from E5-E7. Stagia3 plasmid encoding AP and GFP was used to assay for CRM activity (purple stain), and was co-electroporated with a control plasmid (CAG-mCherry, red). Corresponding images of each electroporated explant with CRM activity are shown. (C) Sections of chick retinal explants (E5-E7) electroporated with Oc1 CRM A, B, J, K and L are shown using the Stagia3 GFP readout. IHC for Oc1 protein (purple) or Lhx1 (gray), a horizontal cell marker, are shown. Examples of cells with GFP expression (arrows) that also expressed Oc1 and Lhx1 proteins are shown with white arrows, with those that showed Oc1 but not Lhx1 are indicated with red arrows. Cells with GFP expression but no overlap with Oc1 and Lhx1 are shown with yellow arrows.

### Identification of CRMs active in developing cones

We then used the differential ATAC-seq profiles to search for CRMs of other genes enriched in the ThrbCRM1+ population, to identify additional regulatory regions potentially relevant for cone development (Figure 4). Near ATAC-seq peaks identified as differentially high among the ThrbCRM1+ cells relative to the ThrbCRM1− cells, one of the most differentially expressed gene was Rbp4 ^31^. Rbp4 is a plasma carrier of retinol and has been shown to be expressed in the scleral portion of the developing retina ^31,44^. Its importance in retinal function was shown by its deficiency or overexpression, which impair vision and lead to retinal degeneration ^45,46^. Similarly, an RBP4 antagonist, A1120, results in a loss of cones ^47^. We tested four ATAC-seq peaks enriched in the ThrbCRM1+ cells that are near the Rbp4 gene, using the AP reporter assay in retinal explants. Three peaks were positive for AP expression (Figure S3A). Rbp4 CRM A and D led to a strong enrichment of GFP expression in cells of the photoreceptor layer that also labelled with expression of a tdTomato reporter driven by ThrbCRM1 (ThrbCRM1-tdTomato) and IHC for visinin (Figure 4A, S3G).

**Figure 4:**
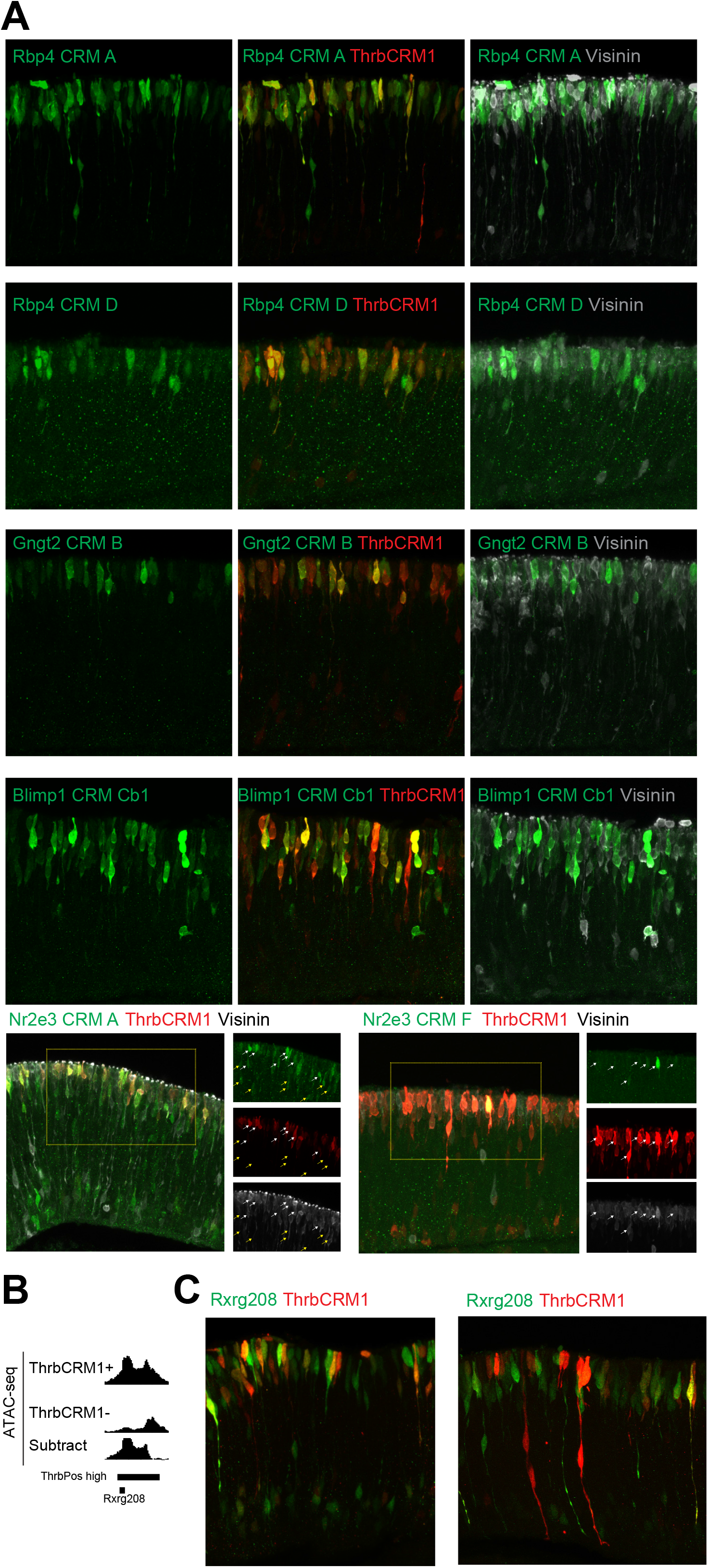
Identification of CRMs active in developing cones nominated by differential ATAC-seq peaks. Assay of CRMs nominated by differential ATAC-seq was carried out by Stagia3 plasmid electroporation of chick explants. Co-electroporation of ThrbCRM1-tdTomato was used to mark developing cones/HCs and their RPCs. (A) Rbp4 CRM A, Rbp4 CRM D, Gngt2 CRM B, Blimp1 CRM Cb1, Nr2e3 CRM A and Nr2e3 CRM F (green) were active in cells with the location and morphology of cone photoreceptors, that expressed ThrbCRM1-tdTomato (red), and expressed the visinin protein (gray). Nr2e3 CRM A showed activity also in cells negative for ThrbCRM1 and visinin (yellow arrows). (B) A differential ATAC-seq peak at the Rxrg locus that overlapped with a previously described CRM for Rxrg, Rxrg208 ^87^, is shown. (C) The CRM activity of Rxrg208 (green) and ThrbCRM1 (red) were compared by co-electroporation of chick explants.

We also studied the regulation of Gngt2, an important component of the phototransduction cascade, an early marker of cone differentiation ^48,49^, and a gene upregulated by addition of thyroid hormone ^50^. We identified two regions more open in ThrbCRM1+ cells that drove AP activity (Figure S3B). A recent study in the mouse retina showed that a reporter for mouse Gngt2 (mGngt2Enh1) showed activity in both cones and rods ^51^. Gngt2 CRM A in the chick showed conservation with mGngt2Enh1. To assess the cellular activity of Gngt2 CRM B, we electroporated the reporter plasmid with the ThrbCRM1-tdTomato reporter. A very strong overlap between the reporters suggested that indeed, the peaks for Gngt2 were active in cells that were cones (Figure 4A).

Blimp1 has been shown to be under the regulation of Otx2 via the CRM B108 in neonatal mouse retina ^52^, where it regulates the choice between photoreceptor and BP cell fate ^53,54^. We identified several differential ATAC-seq peaks at the Blimp1 locus and three CRMs were found to induce AP activity (Figure S3C). The chick Blimp1 CRM A peak was able to drive reporter activity and has conservation with the mouse sequence from B108, which we found is also active in the chick retina (Figure S3C). The Blimp1 CRM C also had activity, which was further refined to a fragment of 420bp, called Blimp1 CRM Cb1 (Figure S3D). We then looked at the cellular activity of the sequence C and its sub-sequence Cb1, co-electroporating the sequences along with the ThrbCRM1− tdTomato reporter. A strong correlation suggested that both elements were active in cones, as suggested by the presence of visinin in these cells (Figure 4A, S3F).

We also identified multiple differential ATAC-seq peaks at the Nr2e3 locus. Nr2e3 is a direct transcriptional target of the rod TF, Nrl in mice ^55^. It has been shown to be required for the repression of a subset of cone genes within rods ^56,57^. It also regulates rod genes in collaboration with Nrl and Crx, although Nr2e3 is not required for rod gene expression ^56–58^. Previous studies showed that Nr2e3 is transiently expressed in cones in zebrafish and in mice ^57,59^, as was also recently reported for chick ^51^. Accordingly, Nr2e3 RNA was enriched in ThrbCRM1+ cells. As ThrbCRM1+ progeny do not become rods ^60^, we wondered if these potential CRMs would be active in cones. We tested seven regions that all showed some level of AP activity (Figure S3E). Two of these elements (Nr2e3 CRM C and D) overlapped with recently described potential CRMs ^51^. Nr2e3 CRM A and F were positive in cones, as shown by their overlap with ThrbCRM1 activity and colocalization with visinin protein (Figure 4A). Nr2e3 A showed broader activity, as it was active in cells negative for ThrbCRM1 and visinin (Figure 4A).

RXRg RNA is also strongly enriched in ThrbCRM1+ cells. Rxrg is a partner of Thrb, and is required for proper cone opsin regulation ^28^. We inspected our data for peaks that were more open in ThrbCRM1+ cells at this locus. We identified a potential CRM near the promoter of Rxrg. An enhancer, Rxrg208, active in chick photoreceptors and HCs, was previously identified within this peak ^61^ (Figure 4B). We then asked if ThrbCRM1 and Rxrg208 were active in the same cells. Stagia3 plasmids with these two CRMs were co-electroporated and found to have a strong overlap in their activity, likely in cones and HCs, as well as their progenitor cells (Figure 4C). While cells rarely showed expression driven only by ThrbCRM1, a large number of cells showed expression driven only by Rxrg208. This pattern suggests that these cells might represent early HCs, which may express Rxrg but not Thrb, and/or different subtypes of cones ^62^.

### Rxrg and Thrb are part of the same regulatory network

As Rxrg and Thrb were previously described to have a collaborative role in cone differentiation and patterning ^28,29^, and their respective CRMs showed activity in an overlapping population, we searched for potential upstream TFs of Rxrg208. TFBSs for both Otx2 and Oc1 were predicted by TRANSFAC within the Rxrg208 sequence (Figure 5A). The predicted Oc1 binding site was very conserved between ThrbCRM1 and Rxrg208 CRMs, with the predicted Otx2 binding site located very close to the predicted Oc1 site in both CRMs. Rxrg expression in cones has been shown to be regulated by Otx2 and the Onecut genes, although it was not investigated regarding whether it was direct ^22,23,63^. To assess the necessity of the Otx2 and Oc1 binding sites within the Rxrg208 CRM, we cloned into Stagia3 the region corresponding to the Rxrg208 sequence with or without Oc1 and Otx2 binding sites (Figure 5B). While the WT fragment showed strong AP activity, the absence of the binding site for either Oc1 or Otx2 led to almost no expression (Figure 5B). The necessity of these sequences was further tested by mutating the core part of the motif for both TFs and these mutants also exhibited dramatically reduced expression (Figure 5B). These results suggest that both Rxrg and Thrb might be regulated by the same upstream TFs, which could facilitate their regulation of cone gene expression and patterning, possibly as heterodimers ^28,29^. Further, the reliance upon these TFs, which are active on ThrbCRM1 in RPCs, suggests that Rxrg208 is active in the same restricted RPCs as ThrbCRM1.

**Figure 5:**
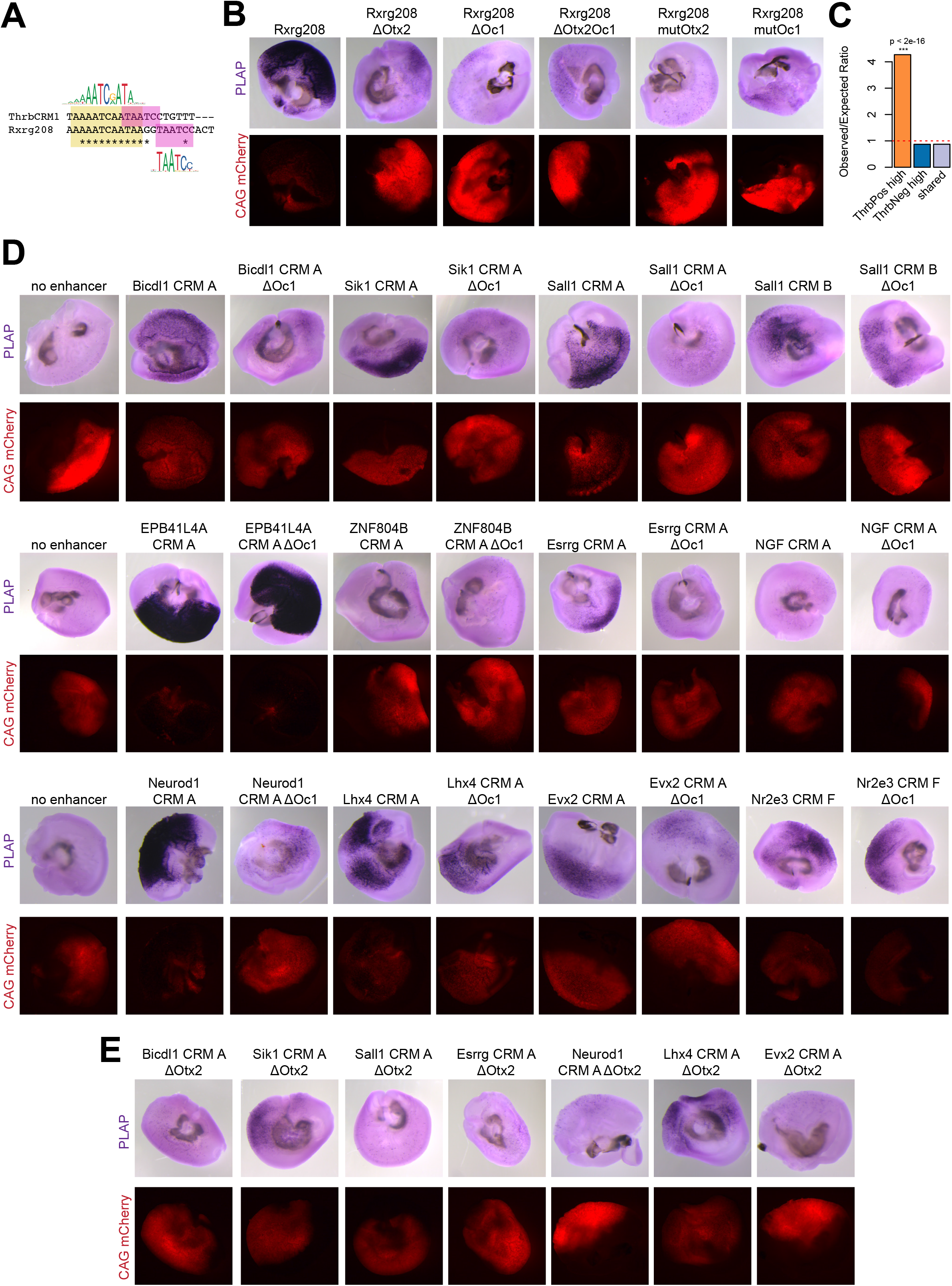
Otx2 and Oc1 regulate multiple genes expressed in developing cones. (A) A comparison of the predicted TFBSs within the CRMs ThrCRM1 and Rxrg208, showed binding sites for Otx2 (pink) and Oc1 (yellow). (B) Deletions and mutations of the Otx2 and Oc1 sites shown in panel A were made and assayed using plasmid electroporation of chick retinal explants. (C) An analysis of the co-occurrence of the TFBSs of Otx2 and Oc1 in ThrbPos high, ThrbNeg high and shared peaks between ThrbCRM1+ and ThrbCRM1− cells. O/E (observed/expected) ratio is calculated to measure the co-occurrence, the p-value is calculated by the Fisher-test (see details in Methods). (D) CRMs nominated by the co-occurrence of Otx2 and Oc1 binding sites within 100 bp were tested for activity using the plasmid electroporation assay. Deletions of the predicted Oc1 sites were made, as indicated, to test for the necessity of this site (E) Regions that showed a dependence on the Oc1 binding site (panel D) were also tested for the necessity of the Otx2 binding site.

### Otx2 and Oc1 regulate multiple other genes of the GRNs for cones

Because the TFBSs for Otx2 and Oc1 were enriched within the ATAC-seq ThrbCRM1+ high peaks (Figure 1D), we asked if the two TFs might have a broader role beyond regulating Thrb and Rxrg. We analyzed the co-occurrence of the binding sites for both Otx2 and Oc1 within the differential ATAC-seq peaks genome-wide, and the distance between them. The co-localization of binding sites for both TFs was more common within the ThrbCRM1+ high peaks compared to the ThrbCRM1-high peaks, or the shared peaks (Figure 5C). Otx2 and Oc1 motifs were usually found within approximately 200 bp, independent of the cell population. However, these two binding sites were located within 50-100 bp from each other when found enriched in the ThrbPos high peaks (Figure S4).

To examine the significance of the Otx2 and Oc1 binding sites found in differential ATAC-seq peaks, we analyzed their co-localization within 100 bp of each other, relative to the closest differentially expressed genes between ThrbCRM1+ and ThrbCRM1− cells, or within a 10 kb or 300 kb distance of their putative targets defined by the BETA analysis (Figure 1C). Three such differential ATAC-seq peaks associated with the most differentially expressed genes included the CRMs ThrbCRM1, Rxrg208, and a predicted CRM for Sik1. Multiple other candidate CRMs were nominated using this analysis (Sup Tables 4). To assess whether these predicted CRMs were active, we tested 12 of these sequences. They were chosen as they were related to retinal development, and/or were associated with differentially expressed genes, and/or had a very short distance between the predicted Otx2 and Oc1 binding sites. We tested the WT sequence and fragments with a deleted Oc1 site, for their ability to drive AP expression (Figure 5D). Ten WT sequences showed AP activity. They varied in their dependence upon the Oc1 binding site with five showing a strong dependence (Bicdl1 CRM A, Sik1 CRM A, Sall1 CRM A, Esrrg CRM A and Neurod1 CRM A), two showing significant, but less dependence (Evx2 CRM A, Lhx4 CRM A), and three had no reliance upon the Oc1 binding site (Sall1 CRM B, Nr2e3 CRM F and Epb41l4a CRM A) (Figure 5D). We tested the effect of deleting the Otx2 binding site for the fragments showing a strong or moderate dependence on the Oc1 binding site (Figure 5E). Compared to the WT sequence (Figure 5D), we observed an almost total loss of AP expression upon deletion of the Otx2 binding site, except for Sik1 CRM A and Lhx4 CRM A, which showed only a moderate decrease.

To ask if Otx2 and Oc1 might regulate the genes identified above, we looked for expression of these potential target genes in cells expressing Otx2 and Oc1. We generated a single-cell RNA (scRNA-seq) profile of the ThrbCRM1+ and ThrbCRM1− cells using the 10X Genomics platform. The transcriptomes of 10,832 ThrbCRM1+ cells and 7,859 ThrbCRM1− cells were recovered. We subdivided the cells based upon expression of Otx2 and Oc1, and found that most of the potential target genes with a reasonable detection rate showed expression in at least some of the cells also expressing both Otx2 and Oc1, relative to those cells that did not express Otx2 and Oc1 (Figure 6A).

**Figure 6:**
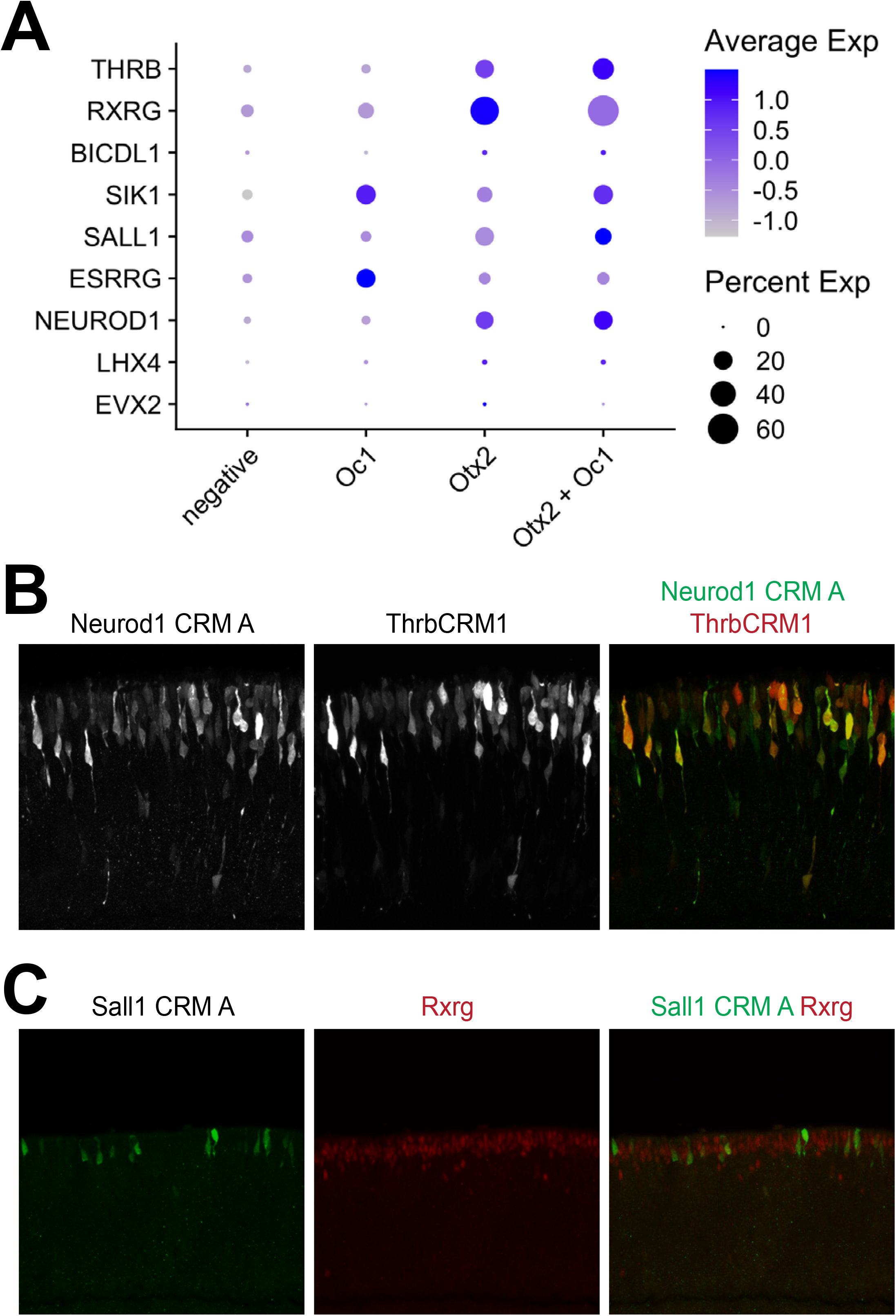
Expression of Otx2 and Oc1 in cells expressing genes with predicted regulation by Otx2 and Oc1. (A) Single-cell RNA-seq was carried out on ThrbCRM1+ and ThrbCRM1− chick retinal cells. The expression of genes predicted to be regulated by Otx2 and Oc1 (Figure 5), along with Otx2 or Oc1 expression, was analyzed, as shown. (B) The Neurod1 CRM A activity (green) was assayed by plasmid electroporation with co-electroporation of ThrbCRM1 (red). (C) The Sal1 CRM A activity (green) was assayed by plasmid electroporation and compared with the expression of the Rxrg protein (red). Binding of Otx2 and Oc1 was evaluated by a CUT&RUN experiment using Otx2 and Oc1 antibodies on whole E5 chick retina.

As Neurod1 is an important regulator of photoreceptor development ^27,64^, and it has a high level of co-expression with Otx2 and Oc1 (Figure 6A), we further analyzed the activity of the Neurod1 CRM A (Figure 5D). We co-electroporated Neurod1 CRM A cloned into Stagia3 with the ThrbCRM1-tdTomato reporter, as a proxy for Otx2 and Oc1 activity. The activity of Neurod1 CRM A showed a strong co-localization with the activity of ThrbCRM1, likely in cones and HCs (Figure 6B). In addition, we electroporated the Sall1 CRM A, as Sall1 was shown to be regulated by thyroid hormone and to be expressed in chick cones ^50,62^. Sall1 CRM A was active in cones, seen by the morphology of the GFP-positive cells and IHC for the Rxrg protein (Figure 6C). Together, these data strongly suggest that Otx2 and Oc1 have a larger role than previously recognized in the regulation of genes in the cone/HC RPC and/or newly postmitotic and differentiating cones and HCs.

### Otx2 and Oc1 proteins co-localize on chromatin throughout the genome

All of the assays described above regarding the regulation of genes relevant for cone development were performed with plasmid reporters. This assay provides for an assessment of the sufficiency of a CRM in an extrachromosomal, contrived setting. However, it cannot provide evidence of direct binding by an implicated TF within the endogenous chromosomal context. In order to test for direct binding by Otx2 and Oc1 on endogenous sites within chick retinal chromatin, the CUT&RUN assay ^65^ was carried out using antibodies against Otx2 and Oc1. Antibodies against the repressive histone mark, H3K27me3, and non-specific IgG were used as positive and negative controls, respectively (Figure S5). We identified 14,293 peaks for Otx2 and 14,246 for Oc1 (Sup Tables 5). To validate the CUT&RUN results, a motif analysis using HOMER was done for both sets of peaks ^66^ (Figure 7). The top predicted motifs for the Otx2 CUT&RUN were for Otx2, GSC and CRX motif (Figure 7A). These three homeobox TFs share very similar motifs for their binding. Interestingly, motifs for Cux2 and HNF6, which are the expected motifs for Oc1, also were found in the most enriched Otx2-bound sequences, further suggesting that Otx2 and Oc1 are often found together. Other top predicted motifs were for other homeobox TFs, Lhx2,3, Nxk6.1 and Isl1, and interestingly, the bHLH genes, Neurod1, Neurog2, Atoh1, Ascl1, Ptf1a and Olig2 (Figure 7A, Sup Tables 6). HOMER *de novo* predicted motifs confirmed an enrichment for motifs similar to Otx, bHLH and CUT genes (Figure 7B). In addition, a motif for nuclear receptors, matching Thrb, was predicted (Sup Tables 6). When looking at the motifs enriched within the DNA sequences bound by Oc1, Cux2 and HNF6 were the top known predicted motifs (Figure 7C). Next were the same homeobox TFs (Lhx1,2,3, Nkx6.1, Isl1) detected as enriched in the sites bound by Otx2, which have very similar binding sites as Otx2 (Figure 7B,C, Sup Tables 6), which again suggests that binding sites for Oc1 and Otx2 are often found together. We could also find enrichment for the bHLHs identified among the Otx2-bound sequences, including Neurod1, which regulates Thrb in mouse ^27^. A major difference between Otx2-bound and Oc1-bound sequences was the higher enrichment for Sox gene motifs for Oc1 (Figure 7B,C). The *de novo* motif prediction for Oc1 binding included those related to CUT, sox, homeoboxes and bHLH genes (Figure 7D, Sup Tables 6).

**Figure 7:**
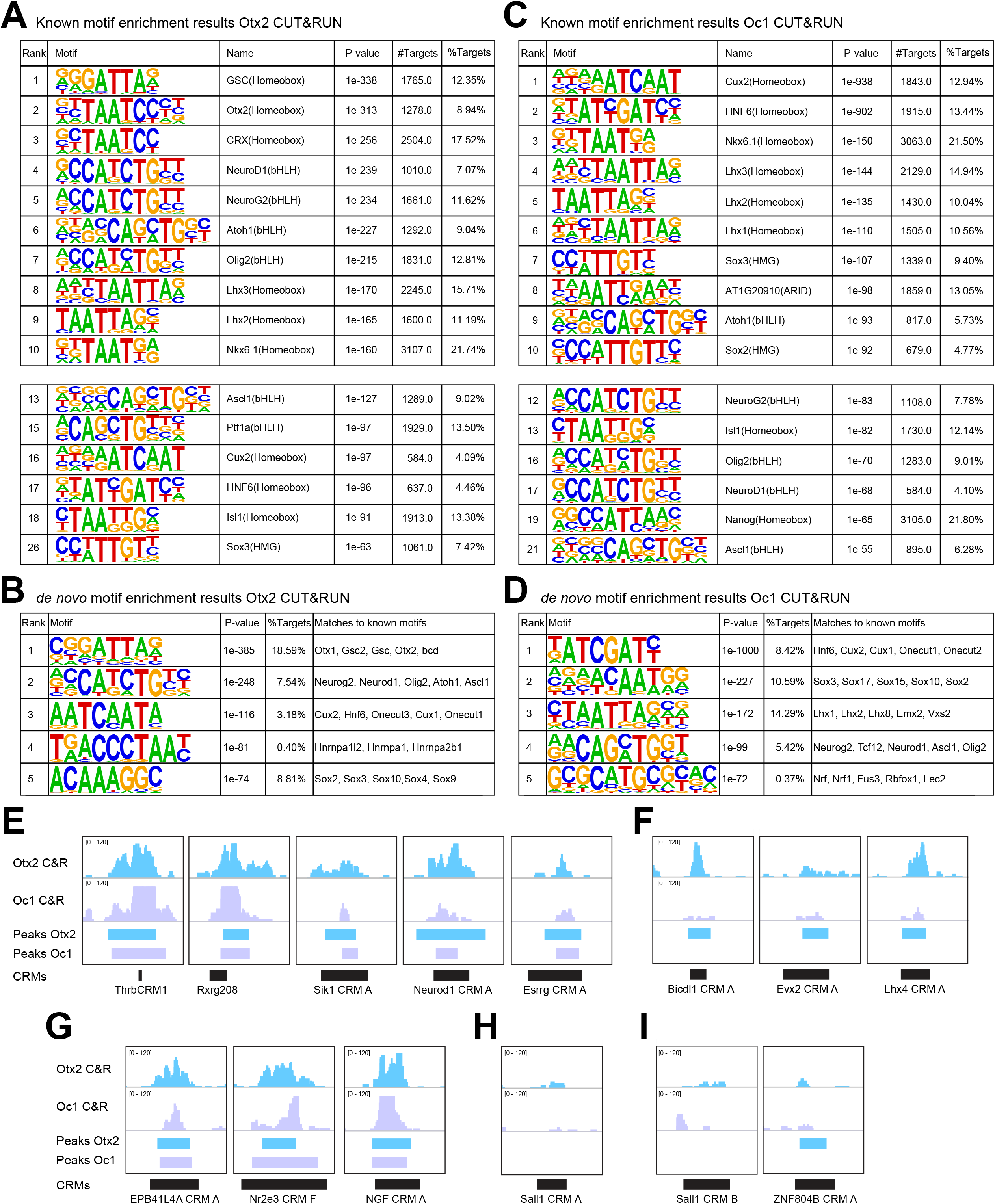
Genome-wide binding of Otx2 and Oc1 proteins assayed by CUT&RUN. (A) The top motifs bound by Otx2 were identified using HOMER, and were for Otx2, GSC and CRX, all homeoboxes related to Otx2. Other frequent motifs were for other homeobox TFs, and for the bHLH TFs, Neurod1, Neurog2 and Olig2. Motifs for Oc1 (Cux, HNF6) were also found to be enriched. (B) An analysis for *de novo* motif enrichment gave results that matched the known motifs for Otx, bHLH and Cut TFs predicted by HOMER. (C) A HOMER analysis of sequences bound by Oc1 showed enrichment for Cux2 and HNF6 TFs, the known motifs for Oc1. Homeobox and bHLH motifs bound by Otx2 also were identified as being bound by Oc1, along with Sox motifs. (D) An analysis for *de novo* motif enrichment gave results that matched the known Cux2 and HNF6 predicted by HOMER. (E-F) CUT&RUN (C&R) profiles were analyzed for Otx2 (light blue) and Oc1 (purple) binding within the CRMs identified as new targets of Otx2 and Oc1 (Figure 5). Peaks called using MACS2 are shown with boxes under the peaks. ThrbCRM1, Rxrg208, Sik1 CRM A, Neurod1 CRM A and Esrrg CRM A showed binding for both TFs (E), while Bicdl1 CRM A, Evx2 CRM A and Lhx4 CRM A showed binding of only Otx2 (F), although deletion of the Oc1 binding site showed a decrease of AP activity (Figure 5). (G) CRMs that did not show a strong decrease in AP staining upon deletion of the Oc1 BS but were bound by both TFs. (H) The Sall1 CRM A had activity dependent upon Otx2 and Oc1 binding sites but was not found to be bound by these TFs using CUT&RUN. (I) Sall1 CRM B and ZNF804B CRM A did not show reliance on the Oc1 binding site for activity and were not bound by Oc1.

As predicted from the motif analysis above, Otx2 and Oc1 CUT&RUN peaks showed a very large overlap within the genome (6,274 peaks) (Sup Tables 5). The list of sites identified as being bound by both Otx2 and Oc1 using CUT&RUN (Sup Tables 5) was compared to the list of CRMs suggested to be regulated by these TFs (Sup Tables 4). About 1/3 of them showed actual binding of Otx2 and Oc1. Interestingly, there were several patterns observed when the CUT&RUN binding data and the results from deletion of the binding sites for Otx2 and Oc1 from the CRM assay (Figure 5) were compared (Figure 7E-I, Table 1). The sequence of the ThrbCRM1 element, which is dependent upon both the Otx2 and Oc1 binding sites, showed binding by both Otx2 and Oc1, as shown previously ^22^ (Figure 7E). Otx2 and Oc1 binding also was seen for the CRMs for Rxrg, Sik1, Neurod1 and Esrrg, all of which showed a loss of activity upon deletion of the Otx2 and Oc1 binding sites (Figure 5B,D,E, 7E). However, some CRMs (near Bicdl1, Evx2, Lhx4) that partially lost activity upon deletion of the Oc1 binding site, showed no significant enrichment for binding by Oc1, though they did show Otx2 binding (Figure 7F). The EPB41L4A CRM A, Nr2e3 CRM F and NGF CRM A had a different pattern. They showed no change in AP activity after the deletion of the Oc1 binding site, but were bound by both TFs (Figure 5D, 7G). An additional case was shown by the Sall1 CRM A. Although the deletion of Otx2 and Oc1 predicted binding sites for these factors led to a loss of activity (Figure 5D,E), the binding of these two TFs was not significantly enriched using CUT&RUN (Figure 7H). Other CRMs, such as Sall1 CRM B and Znf804b CRM A, that did not show a major change in AP activity when the Oc1 binding site was deleted, also, predictably, did not show binding by Oc1 (Figure 7I).

**Table 1.**
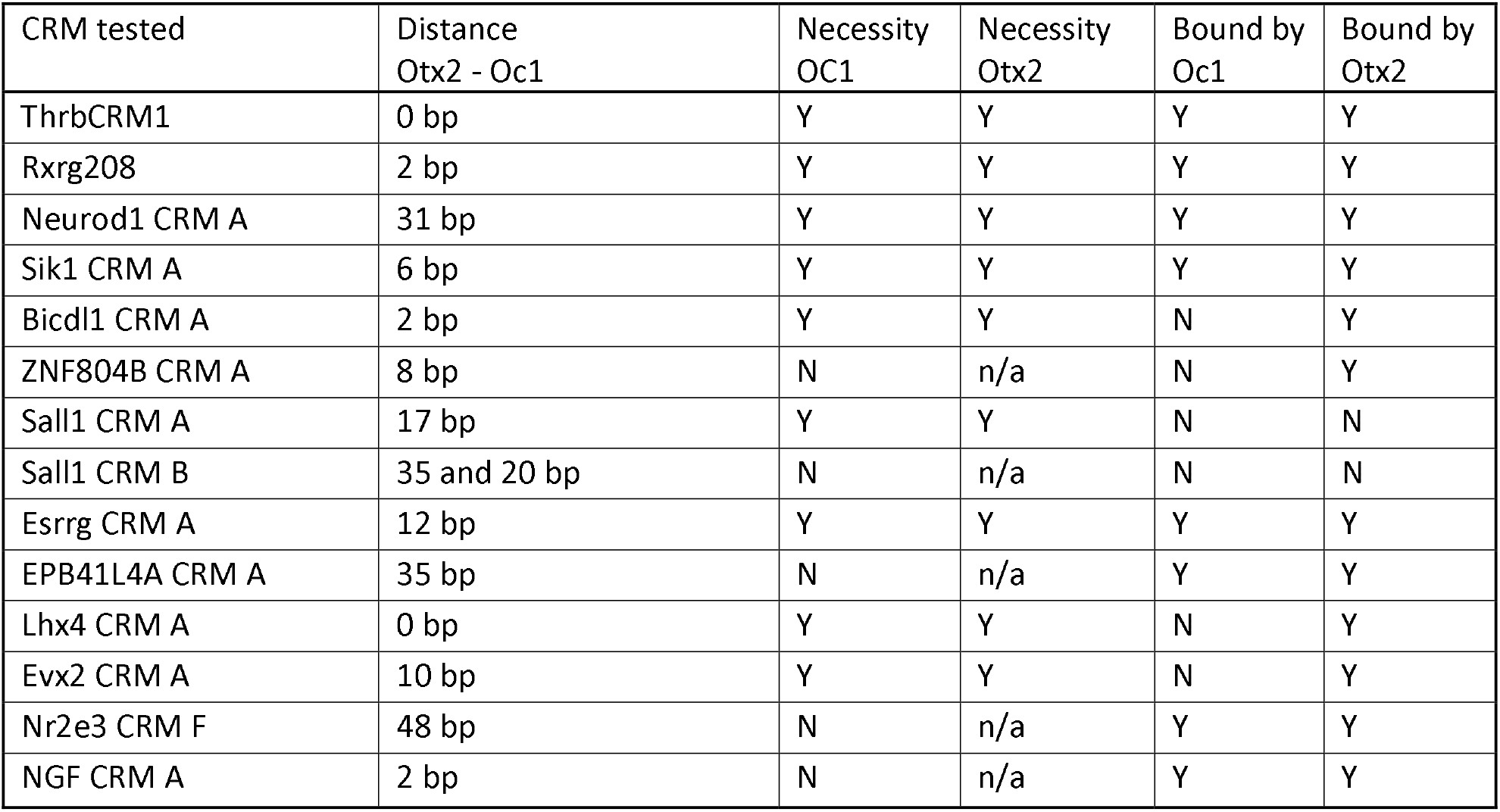
Summary of CRMs tested for direct regulation by Otx2 and/or Oc1. For each CRM nominated by the co-occurrence of the predicted Otx2 and Oc1 binding sites within 100 bp and tested for AP activity (Figure 5), the distance between the binding sites, the necessity of these binding sites, as well as the enrichment of Otx2 and Oc1 binding as measured by CUT&RUN are shown. Each CRM is named for the inferred target gene from the BETA analysis.

The genome-wide CUT&RUN assay for Otx2 and Oc1 binding expanded considerably our previous list of candidate CRMs with co-occurrence of the TFs, defined by known motifs for Otx2 and Oc1. When intersecting the binding data with the ATAC-seq ThrbPos high or ThrbNeg high peaks, we identified several additional potential CRMs regulated by Otx2 and Oc1 at the Thrb, Rxrg, Otx2, Oc1, Sik1, Neurod1 or Sall1 loci, as well as hundreds of other potential CRMs for other genes. The CUT&RUN binding site data also were inspected for the CRMs shown in Figures 2-4 (Figure S6). All of the CRMs at the Otx2 locus, except for Otx2 CRM G, showed enrichment for Otx2 binding. This included Otx2 CRM E and F, further suggesting that Otx2 auto-regulates its expression (Figure 2D,E, S6A)^40^. Some CRMs for Oc1 also showed Oc1 binding, consistent with auto-regulation (Figure S6B). CRMs for Rbp4 were bound by Otx2 but not Oc1 (Figure S6C). Similarly, the Gngt2 CRM A and B showed Otx2 binding only (Figure S6D). The Blimp1 CRM A and Nr2e3 CRM F were bound by both Otx2 and Oc1 (Figure S6E,F).

## Discussion

In this study, we explored the molecular underpinnings of the formation of two early retinal cell types, with a focus on cone photoreceptors, a vulnerable cell type that is critical for color vision. We comprehensively examined the chromatin landscape relative to differential RNA expression, as well as tested many potential CRMs, in cone/HC RPCs and their newly postmitotic progeny. We identified critical sequences within these CRMs by making mutations of predicted TF binding motifs. Binding by these TFs was further confirmed by genome-wide chromatin binding assays. The data from these unbiased approaches show that Otx2 and Oc1 directly regulate a broad range of genes expressed in these early cell types, in many cases acting together on sites that are located relatively close together. These data, along with expression analyses and functional tests in human stem cell cultures ^67^, as well as in mice and chicks ^24–26,68^, indicate that Otx2 and Oc1 are the key TFs for the development of cones and HCs.

The integration of the ATAC-seq data gathered from targeted populations of retinal cells provided a genome-wide view of chromatin in restricted RPCs that were producing cones and HCs. It was interesting to see that the chromatin profiles of ThrbCRM1+ and ThrbCRM1− cells were very similar. The ThrbCRM1− population is presumed to be composed primarily of RPCs and their newly postmitotic daughters, as electroporation preferentially targets these cell types ^30^. A fraction of these cells, however, might become ThrbCRM1+, and thus may show some aspects of chromatin structure that overlap with those of ThrbCRM1+ cells. In addition, they might not have activated the ThrbCRM1 reporter at a high enough level to be selected by FACS. High level expression from this reporter appears to require a multimerized CRM1 sequence ^69^, which we have noted sometimes suffers deletions during growth of the plasmid in bacteria. The opposite pattern, with peaks that are high in ThrbCRM1− cells also open in ThrbCRM1+ cells, could be due to a lag in the closing of chromatin in certain areas only after cells start to differentiate. Analysis of later time points could address this possibility.

Despite the potentially imperfect separation of ThrbCRM1+ and ThrbCRM1− cells, the integration of the differential open chromatin peaks identified by ATAC-seq pointed to potential CRMs for genes preferentially expressed in the ThrbCRM1+ population. This included several new CRMs for Otx2 and Oc1 that complement our recent analyses of their regulation in the mouse retina, where we used non-coding RNA and chromatin profiling to nominate CRMs ^39,40^. The ATAC-seq differential peaks also provided for a prediction of TFBS motifs enriched in these CRMs, with experimental tests of activity validating these predictions in most cases. These series of experiments led to the appreciation of a larger role for Otx2 and Oc1 as direct regulators of genes involved in early retinal development. Analysis of the distance between the binding sites for these two TFs in the ThrbCRM1+ population showed that they were often closer together than one would predict from random, particularly when compared to their distance apart within peaks near genes that were not preferentially expressed in ThrbCRM1+ cells.

To directly assess the binding of Otx2 and Oc1, a genome-wide CUT&RUN was used. This assay showed that the two TFs indeed often bind within the same regulatory regions. Direct binding of both TFs was shown for multiple genes important in cone development, including Thrb, Sall1, Neurod1 and Rxrg. Interestingly, Rxrg is a well described partner of Thrb, as the two proteins establish cone opsin patterning, possibly as heterodimers ^28,29^. We showed that a previously described CRM, Rxrg208, is active in terminally dividing RPCs expressing Thrb. ThrbCRM1 and Rxrg208 were shown to share a very conserved DNA sequence with nearly overlapping binding sites for Otx2 and Oc1 necessary for their CRM activity. In addition to these two regulators of cone development, Neurod1 was found to be regulated by Otx2 and Oc1. Neurod1 is required for the maintenance and expression of Thrb in cones ^27^ and for survival of photoreceptors in mice ^64^. Furthermore, the motif for Neurod1 binding showed enrichment within the regions bound by Otx2 and Oc1, genome-wide, suggesting that Neurod1 likely plays a broader role than was previously appreciated, in collaboration with these homeobox TFs. Interestingly, Sall1 has been proposed to be regulated by thyroid hormone ^50^ and another member of the Spalt family transcription factor, Sall3, regulates HC and cone development in mice ^70^.

One additional gene identified as regulated by Otx2 and Oc1 is Sik1, which was recently reported to regulate photoreceptor development via the Notch pathway in Drosophila ^71^. As Notch is the most upstream regulator of photoreceptor development known in vertebrates ^72–74^, genes regulated by Notch are of interest. Notch represses photoreceptor fates, perhaps through upregulation of Hes genes, which can repress TFs important for neuronal fates ^75^. In keeping with this, an Otx2 CRM (EELPOT), described in mice, is predicted to be under negative regulation by Hes TFs ^76^. Negative regulation of bHLH factors by Notch, in part through Hes TFs, likely limits neuronal fates as bHLH TFs positively regulate many retinal cell fates ^77^. If these Notch-dependent activities lead to the initiation of Otx2 and Oc1 transcription ^41,76^, which then reach a threshold level, they might remain stably expressed via auto-regulation.

The possibility of auto-regulation by both Otx2 and Oc1 is supported by our CUT &RUN data. We found that Otx2 can bind to several CRMs near the Otx2 gene, and Oc1 similarly can bind to regions near Oc1. Such auto-regulation may enable the patterns observed as cells differentiate, where Otx2 stays on and Oc1 goes off in cones, and vice versa in HCs ^24,43,78^. There is likely a role for negative regulation as well, but this has not been explored. Otx2 auto-regulation appears to be via the CRM E in early chick retina, as we report here, and via Otx2 CRM 05 in mouse BP cells, as we found recently ^40^, with these two CRMs sharing highly conserved sequences. Otx2 auto-regulation may be a theme for this gene, as it has been proposed in other contexts as well ^79,80^. Otx2 auto-regulation, as well as its direct binding to cone genes, might suggest that Otx2 is a terminal selector gene ^81^ for photoreceptors. This designation is somewhat ambiguous, however, as Otx2 is also expressed at a high level in BP cells and is required for their development.

Otx2 had been shown to be an important TF not only in the retina, but in several other tissues, where it was found to interact with different partners that modulate its role as a repressor or an activator. It addition to its role in directly regulating transcription, it has been proposed to act as a pioneer factor, regulating access to CRMs within chromatin, notably through collaboration with Neurod1 ^82,83^. Similarly, Onecut TFs have been proposed to act as pioneer factors, as their misexpression led to the generation of neurons from fibroblasts ^84^, as has been shown to follow overexpression of Neurog2 ^85^. The similar phenotypes of neuron induction following Oc1 and Neurog2, and the collaboration of Otx2 with Neurod1, are interesting findings in light of our CUT&RUN motif analysis, where binding sites for combinations of these TFs are overrepresented. Moreover, the notion of pioneer activity of Otx2 and Onecut factors fits in well with their roles in retinal development. Since Onecut genes are not expressed in cones, and Otx2 is not expressed in HCs, yet mouse KO’s show a role for both factors in cone and HC development ^23,24,26^, an early role, perhaps as pioneer factors in RPCs, is supported. As cells exit mitosis to differentiate, the roles of these two TFs may change, and may include some cross-repression ^63^, as well as positive auto-regulation.

Diseases affecting cone photoreceptors, such as Retinitis Pigmentosa and AMD, affect the lives of millions of people. The data presented here bring a greater understanding of the processes that might lead to the development of cell-based therapies to treat retinal disease. In particular, human iPSC-derived organoids produce few cones, especially relative to rod production ^18,67^. The engraftment of stem-cell derived cones, the generation of cones from endogenous stem cells, and/or the use of cones for in vitro screens of potential therapies, would all benefit from an efficient production of cones. Cones are also the target for gene therapy and vectors that direct expression specifically to cones are beneficial for this approach. The CRMs reported here likely will expand the repertoire of CRMs for cone expression, in terms of levels, timing, and/or specificity. Furthermore, there is quite a bit of heterogeneity among individuals with the same retinal disease gene. Some of this variability may result from differences in the CRMs for the disease gene. In addition, many individuals with retinal disease do not have a known disease gene allele. As multiple genes characterized in our study have been associated with retinal diseases, their associated CRMs, or the motifs identified here, can form the basis of a study for causal variants that might affect the expression of a nearby disease gene ^86^.

## Acknowledgments

We thank Bess Miller for her help with the Blimp1 CRM screening, as well as the Microscopy Resources on the North Quad (MicRoN) core, the Department of Immunology’s Flow Cytometry Facility, the BWH Flow Cytometry and the Biopolymers Facility cores at Harvard Medical School for their service and support. We thank Nikki Kong for her generous gift of Protein A-MNase. We thank the Cepko and Tabin lab members for valuable discussions and advice, and for sharing reagents. Support was provided by fellowships from the Swiss National Science Foundation (SNSF) and the Human Frontier Science Program (HFSP) (to N.L) and the National Institutes of Health (NIH) (C.C, N.L, C.L) and Howard Hughes Medical Institute (HHMI) (C.C).

## Author Contributions

N.L and C.C designed the experiment. N.L, C.L and S.W designed the analyses. N.L conducted the experiments, with help from J.C. S.W did the ATAC-seq, RNA-seq integrations and analyses. C.L performed the scRNA-seq analysis. N.L did the CUT&RUN analyses. C.C and P.P supervised the project. N.L and C.C wrote the manuscript, in consultation with all other authors.

## Methods

### Animal handling

Chick embryos procedures were approved by the Institutional Animal Care and Use Committee at Harvard University.

### Electroporation & AP staining

Embryonic day 5 (E5) chick retinas were dissected and electroporated ex vivo as described previously ^30,88^, with 5×50ms 22.5V pulses and 950ms intervals, using a NEPA21 type II Nepagene electroporator. The electroporation chamber was modified as previously described (Montana et al., 2011). DNA was diluted in PBS with 6ug total for the CRMs plasmids, and 3-3.75ug for the control plasmids, in 60ul. After 2 days in culture in 10% FBS, 45% DMEM, 45% F12 and 100U/ml penicillin, retinas were fixed at RT for 30min in 4% Formaldehyde, washed 3x in PBS. After fixation, at least two biological duplicates we processed for AP staining at RT ^88^, using 20ul/ml of NBT/BCIP solution in NTM pH 9.5, and developed for 4-8 hours, or processed for immunohistochemistry. The coordinates (Gallus gallus genome galgal5) or sequences of the regions tested in this study are found in the Supplementary Tables 7. Plasmids were constructed using gBlock or PCR to clone the CRMs sequences in the Stagia3 backbone ^88^.

### Immunohistochemistry & Imaging

Fixed retina were frozen in 30% sucrose/PBS as described previously ^89^ and 20-30um sections were prepared for staining ^90^. Blocking solution was 0.3% Triton X-100 in 1x PBS. Primary antibodies used in these study were: chicken anti-GFP (1:1000, Abcam, AB13970), rabbit anti-mCherry (1:1000, Abcam, 167453), rabbit anti-Otx2 (1:200-400, Proteintech, 13497-1-AP), mouse anti-Rxrg (1:50, Santa Cruz Biotechnology, sc-365252), rabbit anti-Oc1 (1:100, Santa Cruz Biotechnology, sc-13050), mouse anti-Lhx1 (1:30, DSHB, 4F2-c), mouse anti-Visinin (1:250, DSHB, 7G4), rabbit anti-Blimp1 (1:1000, GenScript, A01647). Retinas were washed PBS, mounted with Fluoromount-G (SouthernBiotech). Retina explants were imaged on a Leica M165FC microscope. Retinal section images were acquired using a Zeiss LSM780 inverted confocal microscope. Images were processed with ImageJ to adjust brightness and contrast.

### FACS sorting

E5 retina were electroporated with ThrbCRM1 and CAG-mCherry plasmids and cultured ex vivo for 2 days. Retina were dissociated as described previously ^91^ using DAPI as a dead cell stain. Cells from retina not electroporated, or electroporated with only ThrbCRM1 or CAG-mCherry were used to determine background. Cells from 2-3 retinas were collected and sorted as ThrbCRM1+/mCherry+ and ThrbCRM1-/mCherry+ using a BD FACS Aria machine, and processed for ATAC-seq or 10X scRNA-seq protocols.

### ATAC-seq

ATAC-seq libraries for each condition were prepared using the standard protocol ^92^. We used ~50,000 cells per condition, from retina cultured 2 days *ex vivo*. The first replicate was performed and sequenced separately. The second and third replicates were composed of cells sorted on the same day from the same pool of dissociated retinas, prepared for ATAC-seq and sequenced in parallel. scRNA-seq

Sorted cells were washed in PBS 0.04% BSA and the single cell suspension were between 600 and 1500 cells/ul for target cell recovery of 2000, 7000 cells for replicate 1,2, for ThrbCRM1+ and ThrbCRM1− samples, respectively. All libraries were prepared after the 10x Genomics Single Cell 3’ Reagent Kits v2.

### CUT&RUN

We performed the CUT&RUN according to the protocol from ^93^ and available at dx.doi.org/10.17504/protocols.io.zcpf2vn. E5 WT retina were dissociated as described previously ^91^ to collect ~1,300,000 cells. After ConA-coated magnetic bead binding to cells, the cells were split in 6 tubes for antibody incubation at 4C overnight. Antibodies used were rabbit anti-H3K27me3 (C36B11) (1:100, Cell Signaling Technology 9733T), rabbit anti-mouse IgG H&L (1:100, ab46540), rabbit anti-Oc1 (1:100, Santa Cruz Biotechnology, sc-13050) and rabbit anti-Otx2 (1:200, Proteintech, 13497-1-AP). The chromatin digestion and release step was following the protocol option 1: Standard CUT&RUN. Each sample was washed in 300ul Dig-wash buffer, then we added 6ul of 100mM CaCl2 for chromatin digestion and release and samples were incubated in cold block on ice. 100ul were collected after 25min and added to a new tube containing 2X STOP Buffer, similarly 100ul were removed after 30min and the remaining 100ul were digested for 45min before stopping digestion and proceeding with the protocol. Libraries for cells digested for 25 and 30 min were then prepared with NEBNext Ultra II DNA Library Prep kit (E7645S), following the protocol available at dx.doi.org/10.17504/protocols.io.wvgfe3w, except for samples treated with IgG and H3K27me3 antibodies that were prepared following NEB protocol, as recommended.

### ATAC-seq data analysis

ATAC-seq data sets were aligned to the Gallus gallus genome galgal5 using bowtie2 ^94^ with -X 1000. MACS2 ^95^ was used to identify the ATAC-seq enriched regions. Parameters -B --SPMR --nomodel -- extsize 146 were used while peak calling. MACS2 bdgcmp (-m subtract) was used to calculate the subtraction between two conditions; Peaks with p-value <= 1e-30 that identified by MACS2 bdgpeakcall based on the subtracted ATAC-seq signal between conditions were defined as the differential peaks. This approach was used as we found that some ThrbPos high peaks partially overlapped ThrbNeg high peaks, and within such regions, differential enrichment in sub-regions might not be called differential by MACS2. Preliminary analyses of CRM activity within differential sub-regions of such peaks showed that some of these regions displayed specific CRM activity.

### RNA-seq data analysis

RNA-seq data sets were aligned to Gallus gallus genome galgal5 using STAR ^96^ with ENCODE standard options. RSEM ^97^ was used to do the transcript quantification, and differential expression analysis were performed with DESeq2 ^98^.

Selected peaks activating and repressive function analysis

We used binding and expression target analysis pipeline ^32^ with parameter --df 0.05 to predict different classes of peaks’ activating and repressive function. Regulatory potential for each gene was calculated as 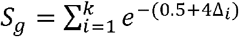. All peaks (k) near the transcription start site (TSS) of the gene (g) within a 100 kb are considered. Δ is the exact distance between a binding site and the TSS proportional to 100 kb (Δ = 0.1 means the exact distance = 10 kb). P-values listed in the top left were calculated by Kolmogorov-Smirnov test to measure the significance of the up-regulated genes group or down-regulated genes group relative to static genes group.

### Motif analysis

MDSeqPos (X. Shirley Liu laboratory) was applied to identify binding motifs based on ThrbPos high peaks and ThrbNeg high peaks. All binding sites were trimmed or extended to 600 bp and centered at the center of the peak regions. Motifs were ranked by the z-score that calculated from MDSeqPos, only top 70 motifs with expression in at least one condition were collected. The list of motifs identified in each condition can be found in Supplementary Tables 3. The enrichment significance in each class were shown in Figure 1D. Values shown in Figure 1D are negative z-score output from MDSeqPos, the higher the value, the more significant the motif enriched, value = 0 represents the lack of enrichment.

### Motif co-occurrence analysis

FIMO (tool from the MEME suite, version 5.0.3, with parameter --thresh 1e-3 --max-stored-scores 2000000) ^99^ was used to search all hits of Otx2 and Oc1 across the whole genome. Bedtools (shuffleBed) ^100^ was used to get the random peaks with the same feature of ATAC-seq peaks from whole genome. Fisher-exact test was used to calculate the p-values between ThrbPos/ThrbNeg high peaks and shuffled peaks.

### CUT&RUN analysis

The reads from the CUT&RUN experiments were aligned to Gallus gallus genome galgal5 using bowtie2 ^94^ with the parameters: --local --very-sensitive-local --no-unal --no-mixed --no-discordant -- phred33 -I 10 -X 700 and processed with SAMtools ^101^ to remove duplicates and MT reads. We generated BigWig coverage tracks using deepTools bamCoverage (Ramírez et al. 2016) with the following parameters: -binSize 1 --normalizeUsing RPGC --effectiveGenomeSize 1046932099. BigWig for all datasets were visualized in IGV Genome browser. Peaks were called using MACS2, with the IgG control of the corresponding digestion time as the background, using the parameter -g 1.03e9. Peaks for Otx2 CUT&RUN that were run in parallel with 25 and 30 min digestion were merged, as was done for Oc1 peaks, using bedtools merge command ^100^. To compare how many peaks were bound by both Otx2 and Oc1, we used bedtools intersect with parameter -f 0.5 -F 0.5 -e. Motif analyses of the CUT&RUN merged peaks was performed with HOMER ^66^, using the parameter -size 50 -p 10.

### scRNA-seq analysis

Libraries were sequenced on the Illumina HiSeq or Novaseq platform, using paired-end reads, with the following read length: 26 bp Read 1 for cell barcode and UMI, 8 bp i7 index for sample index, and 98 bp for transcript. Sequencing output was demultiplexed by the Cell Ranger pipeline, and the unique molecular identifier counts (UMI counts) for each gene were derived with Galgal-5.0 Ensembl transcriptome annotation. The number of cells in each sample was estimated by the Cell Ranger software. All downstream analysis used Seurat v3.1.5 analysis pipeline ^102^. Quality control applied to remove cells with high mitochondrial content or high gene content suggestive of doublet cells. Based on the UMI counts, the cells were subdivided into Otx2 only, Oc1 only, Otx2 and Oc1 double positive and double negative cell population. Log normalization was applied to the dataset to derive the average expression of each subsets.

#### Data availability

The datasets generated during this study are available in the GEO database with the accession numbers GSE151948.

#### Supplementary Information

**Supplementary Tables 1: Differential ATAC-seq peaks for ThrbCRM1+ and ThrbCRM1− cells.**

**Supplementary Tables 2: RNA differential expression and BETA analyses.**

**Supplementary Tables 3: Differential ATAC-seq peaks motif analyses.**

**Supplementary Tables 4: Analyses of Otx2 and Oc1 co-localization within 100 bp of each other, relative to differentially expressed genes.**

**Supplementary Tables 5: Otx2 and Oc1 CUT&RUN peaks.**

**Supplementary Tables 6: HOMER motif analyses on Otx2 and Oc1 CUT&RUN peaks.**

**Supplementary Tables 7: Coordinates or sequences of the CRMs tested in this study.**

---

**Figure S1: (related to Figure 2)**

(A) The activity of the Otx2 CRM E and F was subjected to deletion analysis to identify a shorter sequence responsible for their AP activity. A 161bps for Otx2 CRM E, Otx2 CRM E2.1, and a 410bp fragment, Otx2 CRM F2, were identified as sufficient for activity.

(B) AP staining of chick explants electroporated with the Otx2 CRM F sub-regions.

(C) Using TRANSFAC, several TFBSs were predicted within Otx2 CRM E2.1. We deleted several clusters of predicted TFBSs (del1, 2, 3 and 4), including a region corresponding to an Otx2 BS (del3).

**Figure S2: (related to Figure 3)**

We used ATAC-seq to nominate Oc1 CRMs.

(A) Oc1 CRMs having no AP activity are shown.

(B) Cellular resolution of Oc1 CRM G (green), compared to ThrbCRM1 (red) or Lhx1 (grey), a HC marker. Cells positive for Oc1 CRM G were morphologically resembling RGCs.

**Figure S3: (related to Figure 4)**

Chick retinas were electroporated with Stagia3 plasmids with potential CRMs nominated by differential ATAC-seq between ThrbCRM1+ and ThrbCRM1− cells and genes expressed preferentially in ThrbCRM1+ cells.

(A) Four regions near the Rbp4 gene were positive.

(B) Two regions near the Gngt2 gene were positive.

(C) Three regions near the Blimp1 gene, including the mouse B108 CRM, were positive.

(D) A smaller region driving activity, Blimp1 CRM Cb1, was identified for Blimp1 CRM C.

(E) Seven regions near the NR2E3 locus were positive.

(F) GFP expression driven by Blimp1 CRM C was compared to tdTomato driven by ThrbCRM1 (red) and visinin protein detected by IHC (grey).

(G) GFP expression driven by Rbp4 CRM C was compared to visinin protein detected by IHC (grey). Examples of cells with GFP expression are shown with yellow arrows.

**Figure S4: (related to Figure 5)**

The distance between Otx2 and Oc1 binding sites was analyzed genome-wide.

(A) The distance between Otx2 and Oc1 binding sites was shorter within ATAC-seq peaks that were preferential to the ThrbCRM1+ cells. Distances are also shown for peaks preferentially open in the ThrbCRM1− cells, as well as the shared and shuffled peaks.

(B) The distance between Otx2 and Oc1 TFBS within ThrbPos and ThrbNeg high peaks, non-specific, shuffled peaks, and compared to whole genome.

**Figure S5: (related to Figure 6)**

CUT&RUN was performed on whole chick retina with 25 and 30 minutes of digestion, using Otx2 and Oc1 antibodies (light blue and purple, respectively), along with negative control IgG, and positive control H3K27me3 antibody (dark blue).

(A) The enrichment profiles for each condition are shown for the Hoxd locus, where H3K27me3 is strongly enriched, as expected by the repression of this cluster in these cells.

(B) Zoom-in of the same tracks, at the Otx2 locus. Peaks called using MACS2 are shown with boxes under the peaks.

**Figure S6: (related to Figure 7)**

CUT&RUN for Otx2 and Oc1 and their respective peaks called using MACS2 are shown at several loci: Otx2 (A), Oc1 (B), Rbp4 (C), Gngt2 (D), Blimp1 (E) and Nr2e3 (F).

