## Supplementary figures and images for "Otx2 and Oc1 directly regulate the transcriptional program of cone photoreceptor development"

### Sup Figure 1

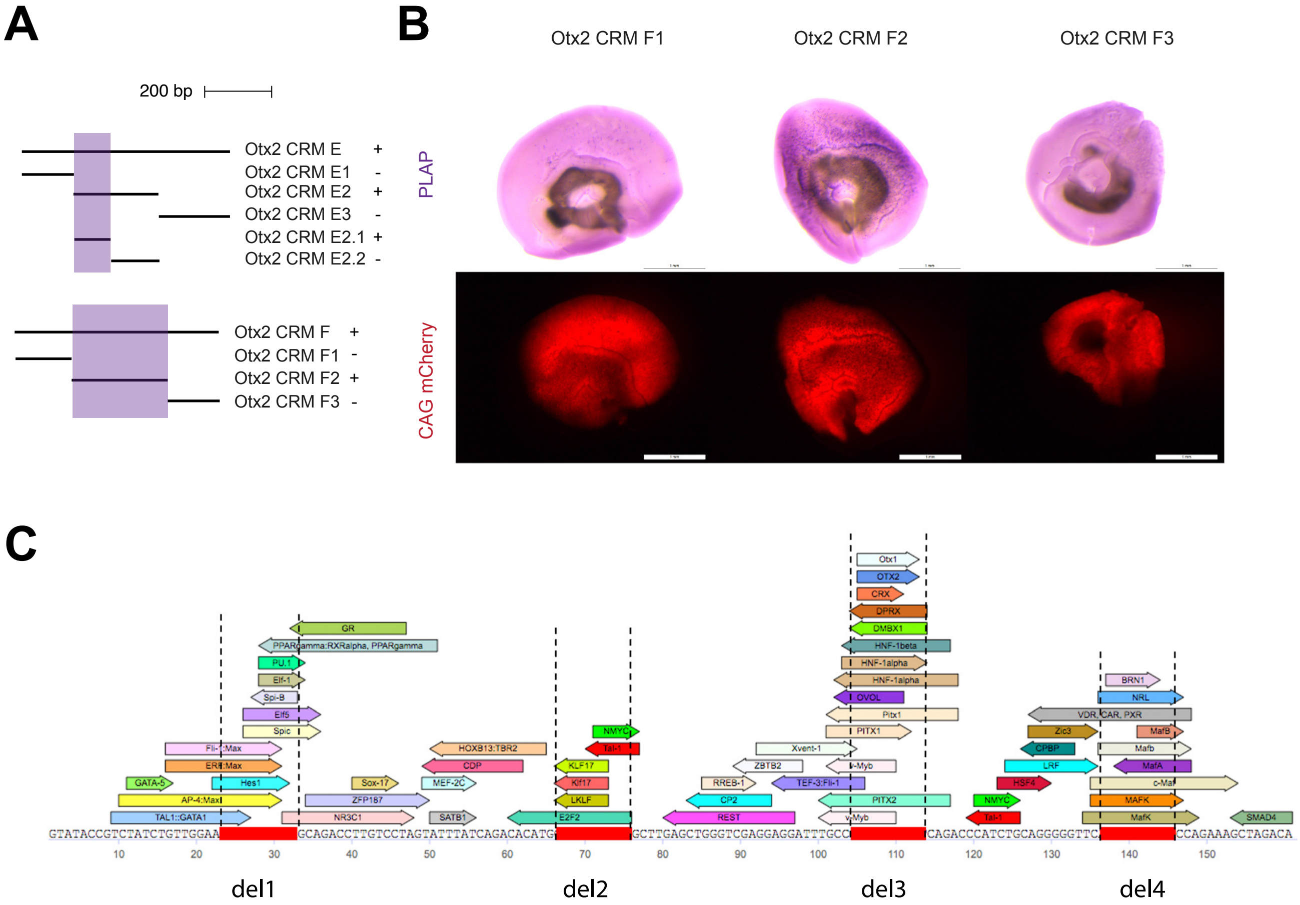

### Sup Figure 2

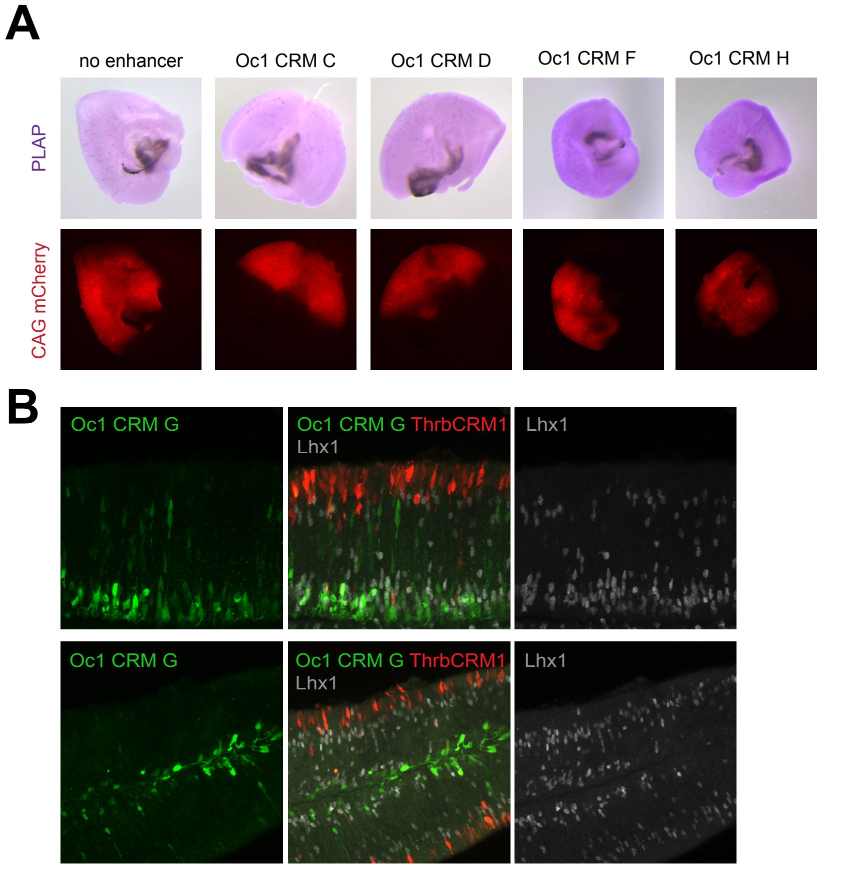

### Sup Figure 3

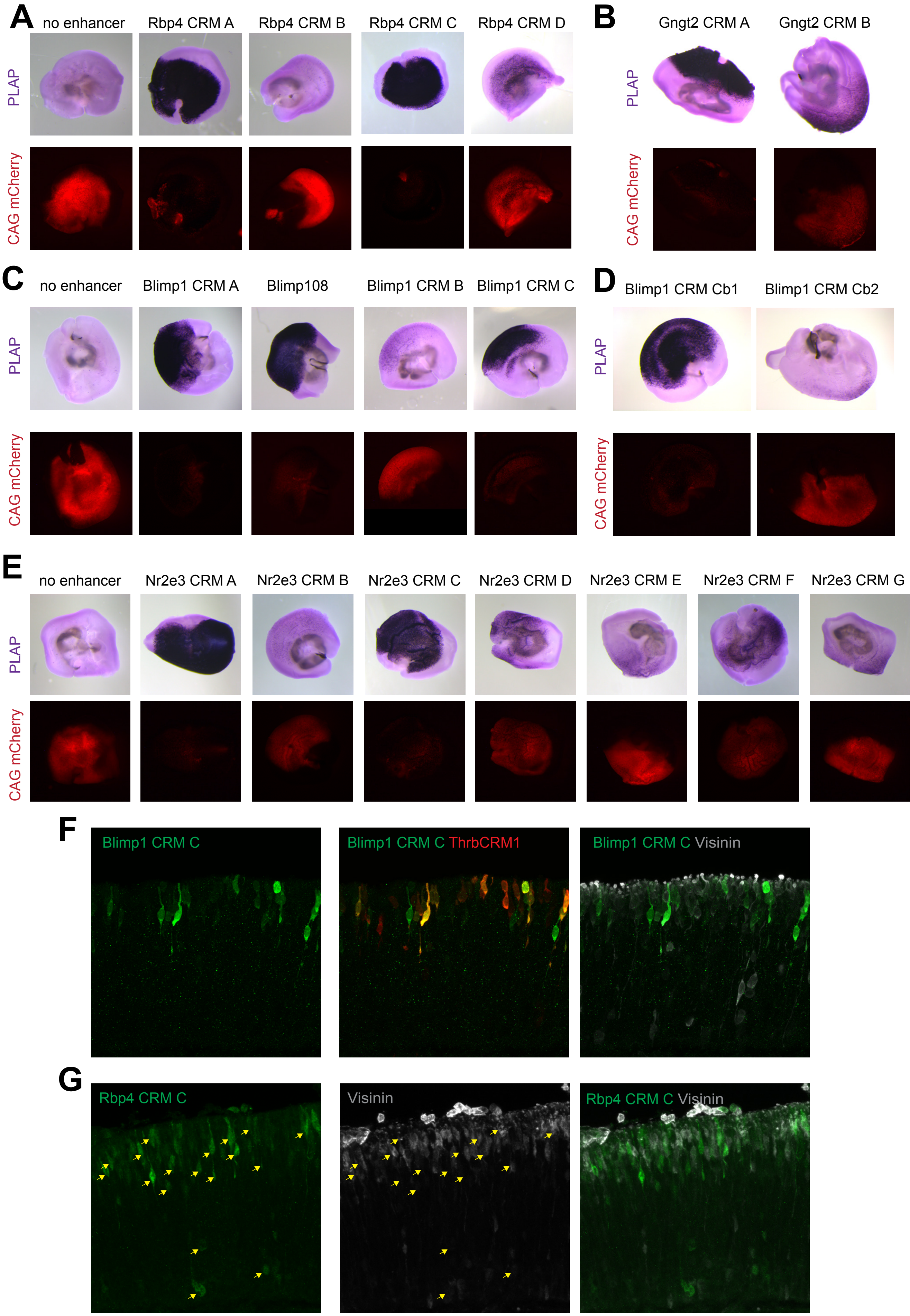

### Sup Figure 4

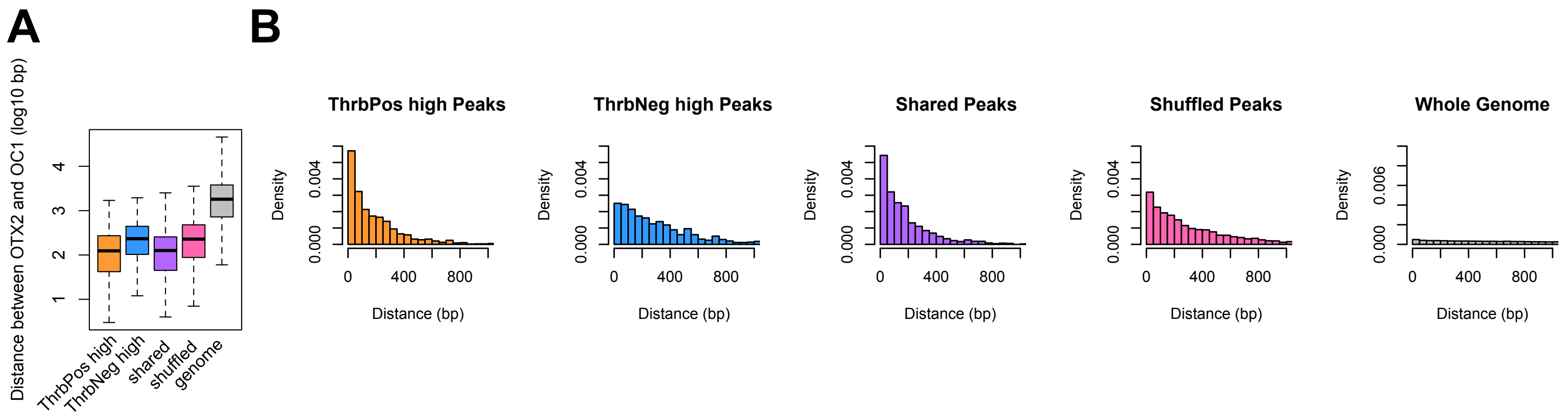

### Sup Figure 5

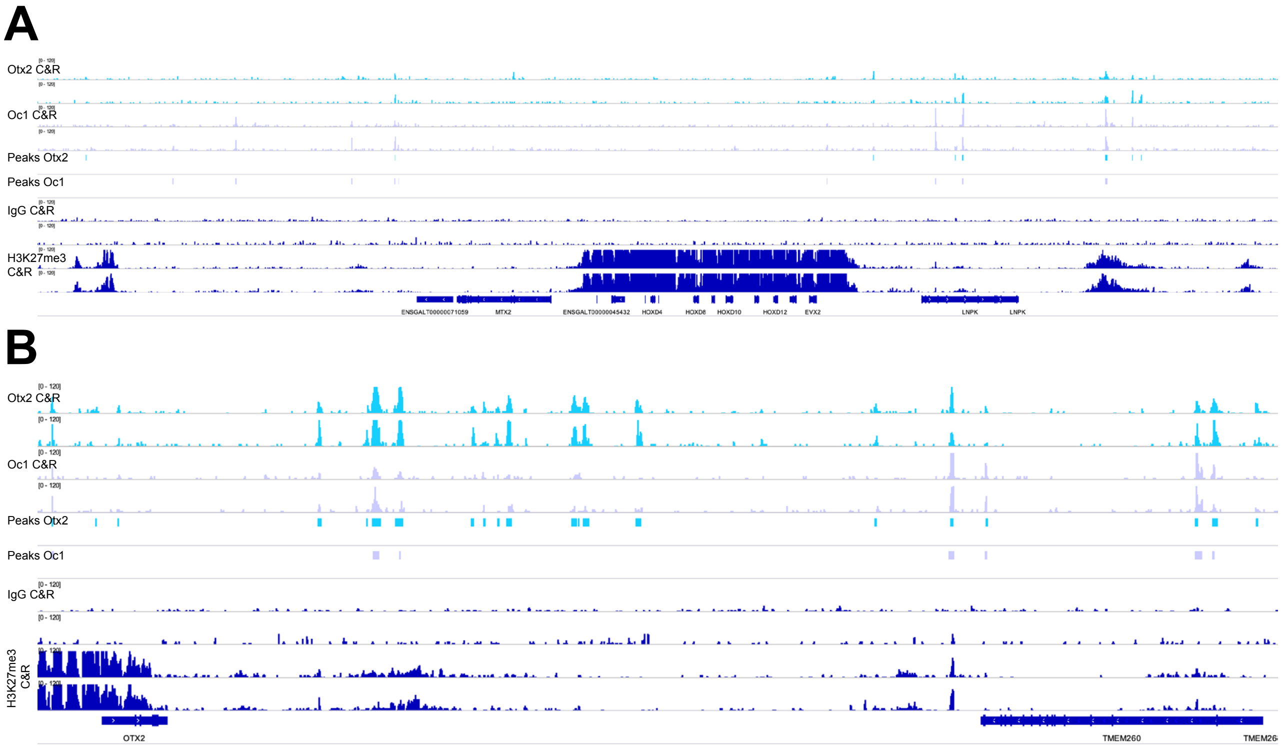

### Sup Figure 6

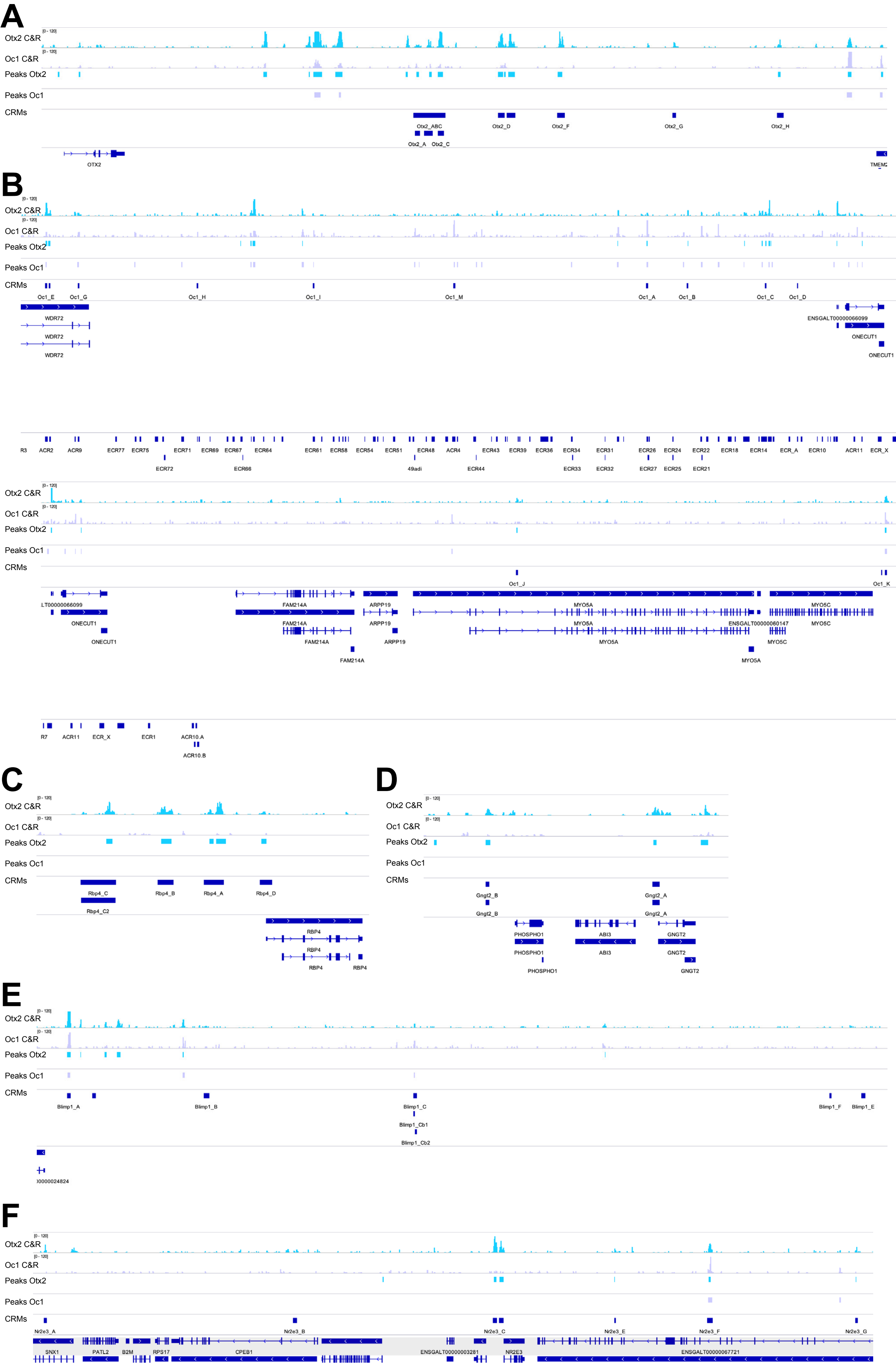
